# Excitatory and inhibitory modulation of septal and striatal neurons during hippocampal sharp-wave ripple events

**DOI:** 10.1101/2020.05.23.112359

**Authors:** Andrew G. Howe, Hugh T. Blair

## Abstract

Single-units were recorded in hippocampus, septum, and striatum while freely behaving rats (n=3) ran trials in a T-maze task, and rested in a holding bucket between trials. During periods of motor inactivity, SWRs triggered excitatory responses from 28% (64/226) and inhibitory responses from 14% (31/226) of septal neurons. By contrast, only 4% (14/378) of striatal neurons were excited and 6% (24/378) were inhibited during SWRs. In both structures, SWR-responsive neurons exhibited greater spike coherence with hippocampal theta rhythm than neurons that did not respond to SWRs. In septum, neurons that were excited by SWRs fired at late phases of the theta cycle, whereas neurons that were inhibited by SWRs fired at early phases of the theta cycle. By contrast, SWR-responsive striatal neurons did not show consistent phase preferences during the theta cycle. A subset of SWR-responsive neurons in septum (55/95) and striatum (26/38) behaved as *speed cells*, with firing rates that were positively or negatively modulated by the rat’s running speed. In both structures, firing rates of most SWR-excited speed cells were positively modulated by running speed, whereas firing rates of most SWR-inhibited speed cells were negatively modulated by running speed. These findings are consistent with a growing body of evidence that SWRs can activate subcortical representations of motor actions in conjunction with hippocampal representations of places and states, which may be important for storing and retrieving values of state-action pairs during reinforcement learning and memory consolidation.

## 1. Introduction

The lateral septum (LS) is a major subcortical output target of hippocampal projection neurons (Raisman 1966). LS sends descending projections to midbrain regions such as the lateral hypothalamic area, substantia nigra, and ventral tegmental area (Risold & Swanson 1997), which in turn send diffuse projections to the ventral and dorsal striatum. It has been proposed that this septal output pathway may be an important route via which the hippocampus exerts influence over behaviors that are regulated by the midbrain dopamine system, including motor actions, reward-seeking, attention, arousal, and decision making (Luo et al., 2011; Gomperts et al., 2015; Tingley & Buzsaki 2018, 2020; Wirtshafter & Wilson 2019). To further investigate how hippocampal output influences the activity of septal and striatal neurons, the present study analyzed how hippocampal EEG states were correlated with single-unit spikes recorded in hippocampus, septum, and striatum.

The rodent hippocampus exhibits distinct patterns of local field potential (LFP) activity during different behavioral states (Vanderwolf 1969). While an animal is actively navigating through its environment, the LFP is synchronized by theta oscillations in the 4-12 Hz band. By contrast, when the animal is at rest, the LFP enters a state of desynchronization punctuated by phasic bursts, a pattern known as large irregular activity (LIA). During LIA, transient synchronization events produce peaks in lower frequency bands (1-50 Hz) of the LFP, known as *sharp waves*. Sharp waves often co-occur with bursts of power in higher bands (125-300 Hz) known as *ripples*. Sharp waves and ripples can occur independently, but they are often observed together in the low and high frequency bands of the LFP (Buzsaki, 2015). Hence, they are commonly referred to together as *sharp-wave ripple* (SWR) events.

Theta and SWR states of the LFP are accompanied by distinct firing patterns of hippocampal neurons. During the theta state, as the animal navigates through its environment, hippocampal *place cells* fire selectively at preferred spatial locations (O’Keefe & Dostrovsky, 1973). Place cells are hypothesized to encode cognitive maps of familiar spatial environments (O’Keefe & Nadel 1978; Redish, 1999), and supporting this, an animal’s momentary position can be accurately decoded from population vectors of place cell activity as it navigates though space (Wilson & McNaughton, 1993). When an animal is at rest, the LFP switches from the theta state to the LIA state, during which SWRs are accompanied by brief population bursts of place cell activity—referred to *compressed replay* events—that can be decoded as “imagined” spatial trajectories through an environment (Skaggs & McNaughton, 1996; Lee & Wilson, 2002; Foster & Wilson 2006; Diba & Buzsaki 2007; Davidson et al., 2009; Karlsson & Frank 2009). While an animal is running on a maze, replay events occur during pauses in motor activity and tend to encode trajectories that start or end at the animal’s current location (Jackson et al., 2006; Johnson and Redish, 2007; Diba & Buzsaki 2007; Karlsson & Frank 2009; Pfeiffer & Foster, 2013; Wu et al., 2017; Kay et al. 2020). These on-maze replay events have been hypothesized to aid in deliberative decision making about where to travel next (Yu & Frank, 2015), and also in assessing outcomes of prior navigational choices (Foster & Wilson, 2006). After an animal is removed from a maze, replay events that occur during subsequent periods of rest often encode trajectories from the recently visited maze environment, and these post-behavioral replay events have been hypothesized to aid in the consolidation of recent experiences to long-term memory (Wilson & McNaughton, 1994; Buzsaki, 1998; Ego-Stengel & Wilson, 2010; Girardeau & Zugaro, 2011).

Here, we analyzed responses of neurons in septum and striatum during SWRs, while freely behaving rats ran trials on a T maze and rested in a bucket between trials. We found that a subset of septal neurons were either excited or inhibited during SWRs; these SWR-responsive septal neurons often fired coherently with hippocampal theta rhythm, and some were modulated by the animal’s running speed, in agreement with other recent reports (Wirtshafter & Wilson 2019). A small percentage of neurons in striatum were also excited or inhibited during SWRs, and as in septum, these neurons spiked coherently with theta rhythm and were commonly modulated by running speed. In both structures, neurons that were excited during SWRs tended to be positively modulated by running speed, whereas neurons that were inhibited during SWRs tended to be negatively modulated by running speed. In septum (but not striatum), SWR-excited neurons fired during late phases of the theta cycle, whereas SWR-inhibited neurons fired during early phases of the theta cycle. After describing these findings in detail, we discuss their possible implications for understanding how the hippocampus modulates subcortical circuits for behavioral learning and decision making.

## 2. Results

Rats (n=3) were trained to run repeated acquisition and reversal trials on a T-maze (Fig. 1). At the start of each session, recording cables were connected and the rat was placed for 5 m in a white plastic bucket located next to the maze (the bucket always remained stationary in the same location, even as the start and goal arms were switched during different trial blocks) for a period of baseline recording. The rat was then placed by the experimenter at the start location on a T-maze apparatus consisting of 4 arms extending 90 cm at right angles from a 30×30 cm central platform (Fig. 1A). Throughout each block of trials, a barrier was placed at the entrance to one of the four arms, while the three remaining arms served as the start, baited, and unbaited arms for the T-maze task (see Methods). After each trial on the maze, the experimenter returned the rat to the bucket for 2-5 m while the maze was cleaned and baited for the next trial.

**Figure 1.**
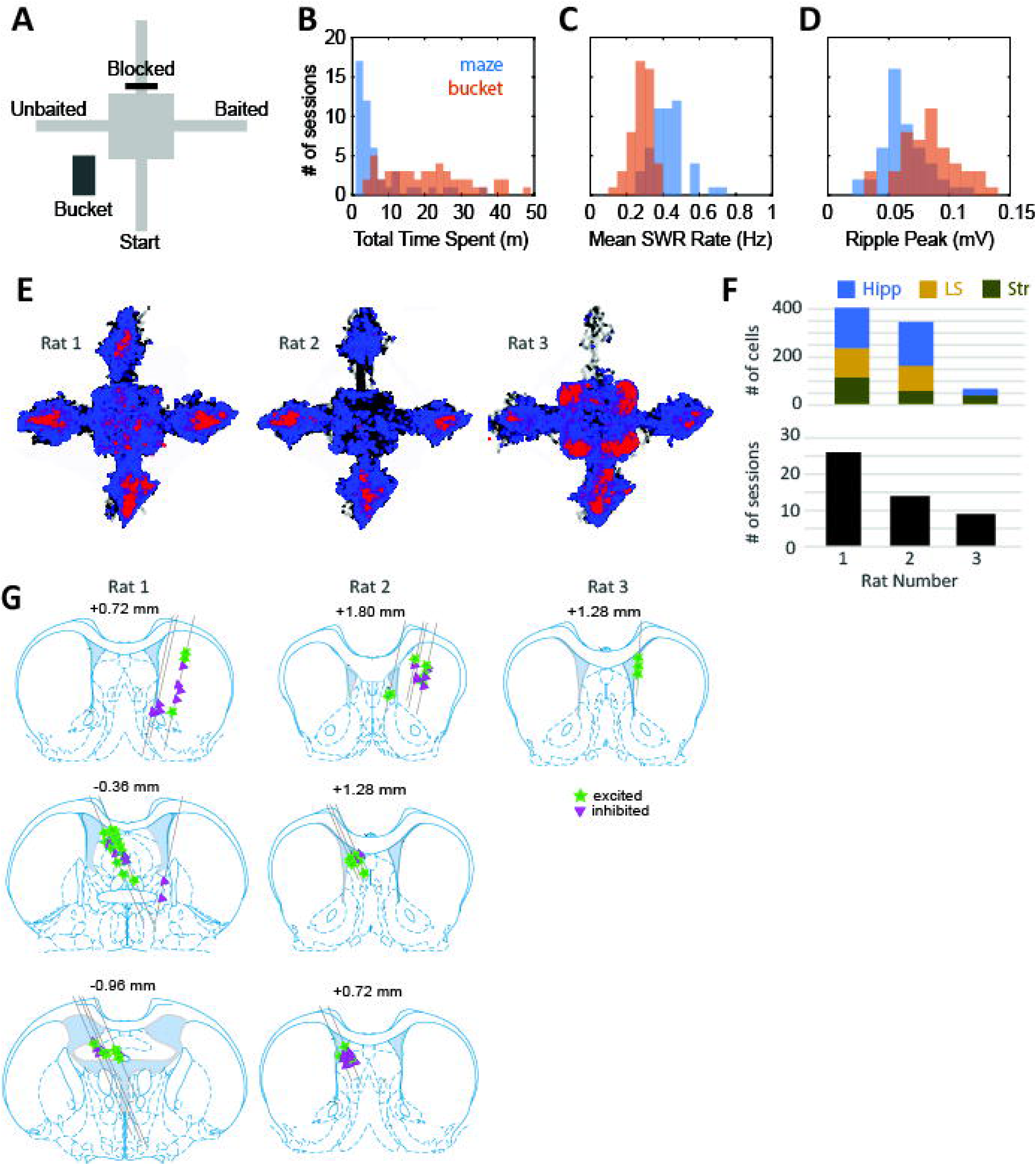
Behavioral and neurophysiological data samples. **A)** Maze apparatus and holding bucket. **B)** Distributions of total time spent sitting still on the maze versus in the bucket during each recording session (N=53). **C)** Distributions of mean SWR rates during stillness on the maze versus in the bucket during each recording session. **D)** Distributions of peak ripple amplitudes during stillness on the maze versus in the bucket during each recording session. **E)** Cumulative spatial distributions across all recording sessions of locations visited (black), locations of sitting still (blue), and locations where SWR events occurred (red) for each rat. **F)** Number of cells recorded in each brain area (top graph) and total number of recording sessions (bottom) for each of the 3 rats in the study. **G)** Septal and striatal recording sites for each rat; symbols indicate recording sites for cells that were excited (stars) versus inhibited (triangles) by SWR events.

Over 6-8 days of initial training, rats learned to find food on one arm of the T-maze. During this initial training period, hippocampal tetrodes were advanced until robust SWRs and theta rhythm were detected on two different tetrodes in the same hemisphere (see Methods). These two tetrodes were assigned as the ripple and theta recording electrodes, respectively, and neither was advanced further during the remainder experiment. Starting with the next session, the goal and/or start arm was changed each time the rat achieved a criterion of 7/8 correct responses (see Methods). Rats spent a median of 3.7 m on the maze and 19.9 m in the bucket during each session (Fig. 1B). In the bucket and on the maze, SWR events were only measured during periods of stillness when the rat’s running speed remained <2 cm/s for 3 s or more (Fig. 1E). During these periods of stillness, the mean rate of SWR generation was significantly higher (paired t_48_=7.97, p=2.4 ×10^−10^) on the maze (0.43 Hz) than in the bucket (0.28 Hz; Fig. 1C), whereas the mean peak amplitude of SWR events was significantly higher (paired t_48_=16.7, p=1.2×10^−21^) in the bucket (84 mV) than on the maze (63 mV; Fig. D).

### 2.1 Unit responses during SWR events

Concurrent with SWR event detection, single units were recorded from the hippocampus (n=216 units from 3 rats), septum (n=226 units from 2 rats), and striatum (n=378 units from 3 rats) during each maze session (Fig. 1F). Hippocampal single units were only analyzed in the hemisphere contralateral from the SWR detection site, to prevent confounds in the analysis that might arise from anatomical proximity between the SWR detection and single-unit recording sites. Hence, in all three brain regions from which single units were recorded, the tetrodes were positioned several mm away from the SWR detection site. To maximize the number of unique cells that were recorded throughout the experiment, tetrodes in septum and striatum were advanced by 333 μm after each behavior session, so that different units would be recorded from these tetrodes in every session. By contrast, hippocampal tetrodes were advanced by at most 83 μm per day (and usually not at all), so that these tetrodes would remain within the hippocampal region throughout the entire experiment. Consequently, most hippocampal units in the dataset were recorded more than once over multiple sessions, whereas each septal and striatal unit was recorded exactly once, during a single session, before the tetrode was advanced to find new cells. Since many hippocampal units were recorded over multiple sessions, the analyses below include only from the first session during which a hippocampal unit was recorded. Thus, in all three structures (septum, striatum, and hippocampus), single-unit responses were always analyzed using a single session’s worth of data for each cell.

Fig. 2 shows example data from Rat 1, obtained from a session during which SWR events were recorded in the right hemisphere of CA1, while hippocampal units were recorded contralaterally in the left hemisphere of CA1 (Fig. 2A). Example recordings are also shown for units recorded in left septum (Fig. 2B) and right striatum (Fig. 2C). Most septal units included in the study were recorded in the lateral septum, but some were recorded the septofimbrial region; few if any were recorded in medial septum (Fig. 1G). A small number of striatal units were recorded from the ventral striatum (nucleus accumbens), but most were recorded in dorsal striatum (Fig. 1G). Hippocampal unit data came from the CA1 region near the pyramidal layer (see examples in Fig. 2A). Single-unit responses to SWR events were measured only during periods of stillness (running speed < 2cm/s), both in the bucket (panels D1-F1) and on the maze (panels D2-F2). By contrast, coherence of single unit spike trains with hippocampal theta rhythm was measured only during periods of active behavior on the maze (running speed >10 cm/s, circular spike phase distributions in panels D3-F3). We adopt the convention that the valley and peak of theta rhythm occur at phases of 0° and 180°, respectively. For the example neurons shown in Fig. 2, SWR events excited CA1 and septum units, but inhibited the unit in striatum.

**Figure 2.**
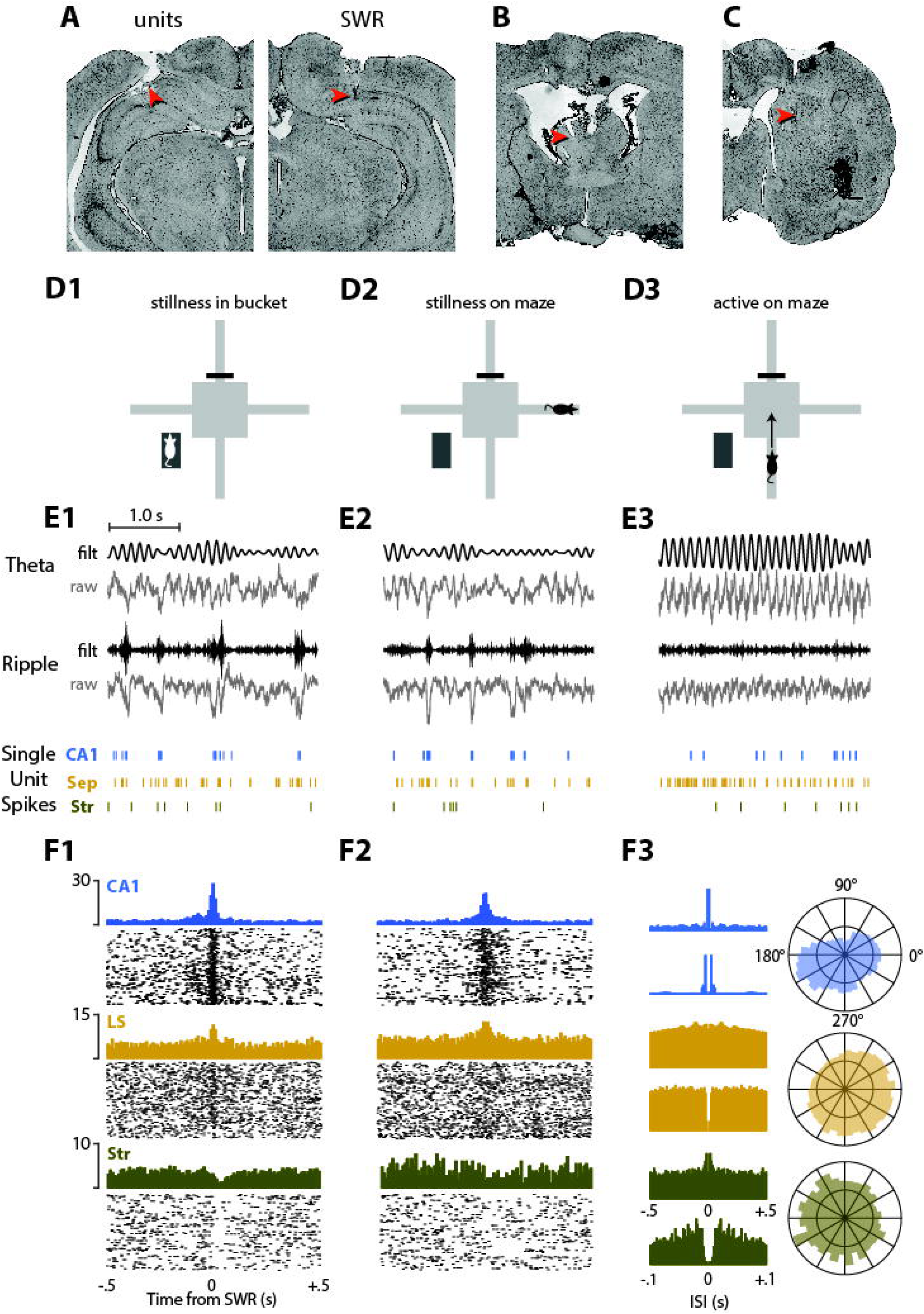
Example data from a single recording session. **A)** Red arrows indicate example unit recording site in the left CA1 (left panel) and SWR detection site in right CA1 (right panel). **B)** Example unit recording site in left septum. **C)** Example unit recording site in right striatum. **D)** Schematic diagrams for three different behavior conditions: stillness in bucket (D1), stillness on maze (D2), and running on maze (D3). **E)** Traces show 3 s of raw and filtered LFP data from the theta (top row) and ripple (middle row) channels, aligned with examples of single unit spike rasters (bottom row) from a CA1, septum (Sep), and striatum (Str); sample data is shown for stillness in bucket (E1), stillness on maze (E2), and running on maze (E3). **F)** Example cell PETHs and rastergrams aligned to SWR events that occurred in the bucket (F1) or on the maze (F2). Autocorrelograms of interspike intervals (ISIs) are shown (F3, left) between -.5 and +5 s (top graph in each row) to illustrate theta rhythmicity, and between and -.1 and +.1 s (bottom graph in each row) to illustrate spike refractory periods. Polar plots (F3, right) show distributions of each example cell’s spike phase relative to hippocampal theta.

#### 2.1.1 Excitatory and inhibitory responses

To test how neurons responded to SWR events, a signed rank test was performed to compare each neuron’s spike rate in a time window spanning ±50 ms from the SWR peak against the baseline firing rate when SWRs were not occurring (see Methods). Two separate tests were performed to analyze whether a neuron responded to SWRs in the bucket versus on the maze, and a neuron was classified as excited or inhibited during SWR events if either of the two tests test yielded p<.01, indicating that the SWR response rate was greater or less than the baseline rate, respectively (Fig. 3). We never observed units with opposing SWR responses for the two behavior conditions (that is, no cells were excited on the maze but inhibited in the bucket, or vice versa). However, some units did exhibit significant SWR excitation or inhibition only in the bucket but not on the maze, or vice versa (see below).

**Figure 3.**
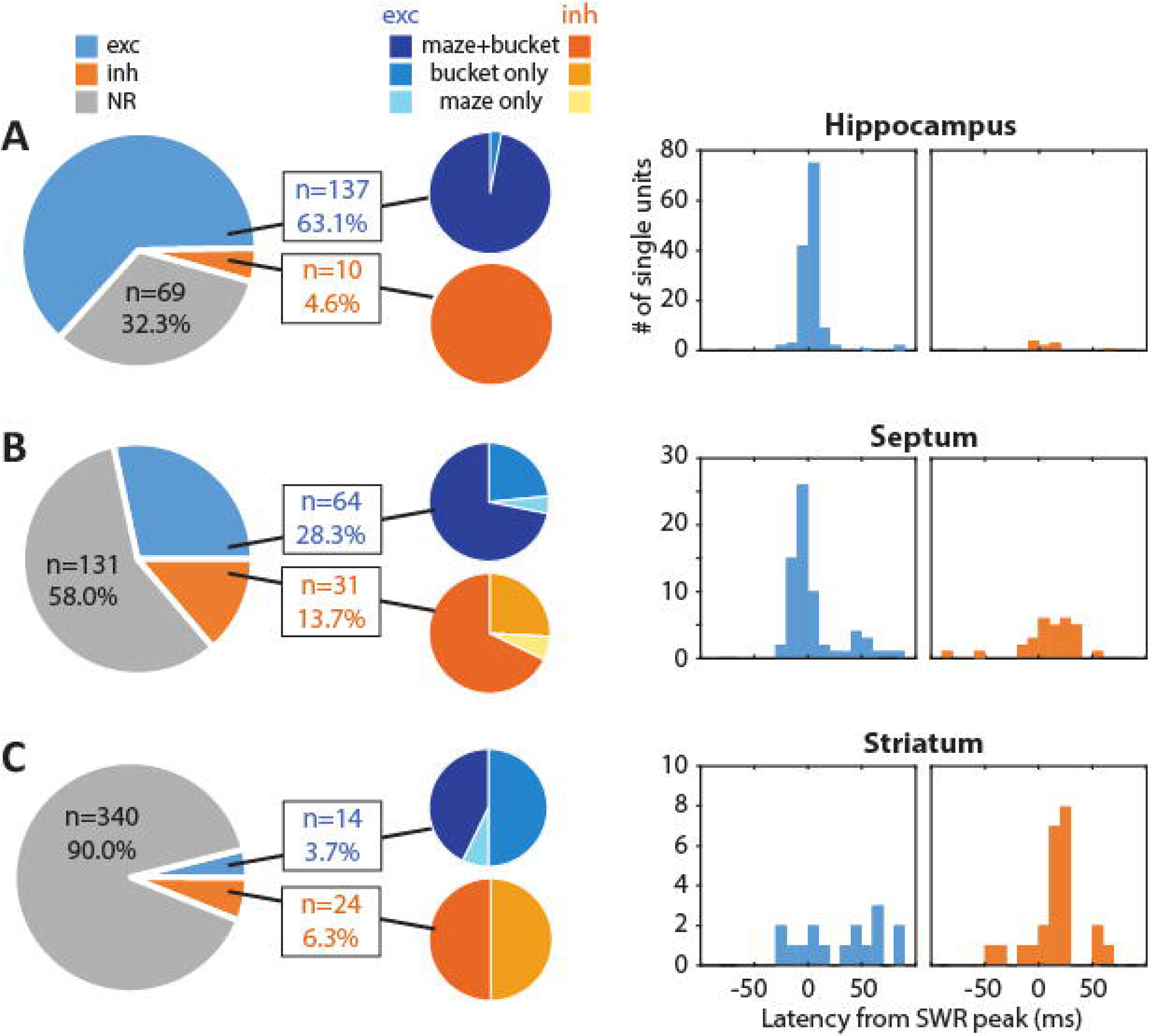
Unit responses to SWR events. Large pie charts in left column show proportions of neurons in hippocampus **(A)**, septum **(B)**, an striatum **(C)** that were excited (exc), inhibited (inh), or non-responsive (NR) to SWR events. Small pie charts in middle column show proportions of SWR-excited (blue) and inhibited (orange) cells that responded in the bucket, on the maze, or both. Histograms at right show distribution of response latencies for cells that were excited or inhibited by SWRs.

##### 2.1.1.1 Hippocampal Units

Hippocampal neurons were recorded from all three rats in the study. Excitatory responses to SWR events were observed in 137/216 (63.4%) of hippocampal neurons, while 69/216 (31.9%) of hippocampal neurons were not significantly responsive during SWR events (Fig. 3A). Only 10/216 (4.7%) hippocampal neurons were inhibited during SWR events, but at least one SWR-inhibited neuron was observed in the hippocampus of each rat. The percentage of hippocampal neurons that were excited during SWRs was similar in all three rats (67.5%, 54.2%, 66.7%). Hippocampal neurons that were excited during SWRs usually exhibited their peak spike response within ±5 ms of the SWR onset (Fig. 3D), with a mean response latency of +4.3 ± 1.4 ms. For this analysis, SWR onset time was measured at the moment when the ripple envelope crossed a standard deviation threshold on the SWR electrode (see Methods). The average latency for inhibitory SWR responses was slightly longer (+10.0 ± 7.1 s), but did not differ significantly from the latency of excitatory responses (t_145_=1.04, p=.3; the statistical power of this comparison was limited by the small sample of SWR-inhibited neurons).

The excitation of hippocampal neurons that we observed during SWRs is consistent with findings from prior studies (Wilson and McNaughton, 1994; Skaggs & McNaughton, 1996; Kudrimoti et al., 1999; Foster & Wilson, 2006; Davidson et al., 2009). Here, we did not decode replay trajectories from hippocampal unit bursts, since this phenomenon has been well studied in prior experiments. Instead, our main objective was to analyze neural activity in septum and striatum during SWRs (see below), and confirming the occurrence of hippocampal unit bursts was necessary to validate accurate isolation of SWR events for these analyses. Note that we only analyze hippocampal unit responses in the hemisphere opposite from where SWRs were recorded. Evidence suggests that SWR events typically originate in CA3, from which they are monosynaptically transmitted to CA1 via Schaffer collaterals (Csicsvari et al 2000; Nakashiba et al., 2009; Buzsaki 2015). If SWR events occur simultaneously in both hemispheres of CA3, then they should arrive nearly simultaneously in both hemispheres of CA1, in which case it would be expected that the peak of the LFP ripple response in CA1 should occur nearly simultaneously with SWR-evoked unit responses in the opposite hemisphere, as we observed.

##### 2.1.1.2 Septum units

Septum neurons were recorded from two of the three rats in the study (n=121 units from rat 1, n=105 units from rat 2). A majority of septum neurons (131/226, or 57.9%) were non-responsive during SWRs, but 64/226 (28.3%) of septal neurons were excited during SWR events (Fig. 3B), which was a significantly smaller proportion of SWR-excited neurons than we observed in the hippocampus, *χ*^2^(1,*N*=442)=54.9, p=1.27×10^−13^, and 31/226 (13.7%) of septal neurons were inhibited during SWR events, which was a significantly higher proportion of inhibited neurons than we observed in the hippocampus, *χ*^2^ (1,*N*=442)=10.8, p=9.9×10^−4^. The overall percentage of SWR-responsive neurons was higher in Rat 1 (39% excited, 30% inhibited) than in Rat 2 (16.2% excited, 18.9% inhibited), possibly owing to the different paths that tetrodes followed through septum in the two rats (see Fig. 1G). The mean latency for excitatory SWR responses in septum was −4.6 ± 3.1 s, which did not differ significantly from the mean latency in hippocampus (t_202_=.07, p=.94). Even though many septum neurons had small negative response latencies, this does not mean that they fired prior to the initiation of hippocampal sharp waves. A small negative response latency simply suggests that the sharp wave signal (which presumably originates from CA3) can be detected a bit earlier by unit spikes in septum than by LFP ripples in CA1. Large negative response latencies (<20 ms) would be more likely to indicate cells that actually fire prior to the initiation of sharp waves, but these were rarely observed. Interestingly, the latency distribution for excitatory septum responses appeared to be bimodal (Fig. 1E): most units responded at short latencies (<10 ms) as would be expected if SWRs originating in CA3 were relayed to septum neurons via a monosynaptic pathway, but some units responded at longer latencies (>35 ms) as might be expected from a polysynaptic pathway (or alternatively, from a slower time constant for integrating excitatory inputs in some septum units). When neurons with long response latencies (>35 ms) were excluded from both the septum and hippocampal datasets, SWR-excited neurons were observed to have significantly shorter response latencies in septum than hippocampus (t_66_=3.65, p=5.2×10^−4^). One possible explanation for this could be that on average, septum neurons might have shorter membrane time constants than hippocampal neurons; if this were the case, then even if SWR events originating in CA3 arrived simultaneously in CA1 and septum, the septum responses could appear slightly earlier than CA1 responses. The mean latency for inhibitory SWR responses in septum was +10.0 ± 5.1 s, which did not differ significantly from the latency of excitatory responses when all excitatory cells were included in the comparison (t_95_=.97, p=.34), but did differ significantly when latencies >35 ms were omitted from both datasets (t_62_=3.16, p=.002). This suggests that excitatory input from hippocampus may monosynaptically drive most of the excitatory SWR-evoked responses in septum, followed at a short delay by polysynaptic feedforward inhibitory responses, and a small number of polysynaptic excitatory responses.

##### 2.1.1.3 Striatal units

Striatal neurons were recorded from all three rats in the study. A majority (340/378, or 90%) of striatal neurons were non-responsive to SWRs (Fig. 3C). Only 14/378 (3.7%) of striatal neurons were excited during SWR events (but at least two SWR-excited neurons were observed in each of the three rats), whereas 24/378 (6.3%) of striatal neurons were inhibited during SWR events (with roughly similar percentages in all three rats: 6.0%, 7.7%, 0%). The latency for excitatory SWR responses in striatum was more variable and on average longer (+36.8 ± 10.8 s) than excitatory responses in hippocampus (t_152_=6.05, p=1.1×10^−8^) or septum (t_82_=4.04, p=1.18×10^−4^; all SWR-excited septum neurons were included in this comparison). The mean latency for inhibitory SWR responses in striatum was +15.0 ± 8.4 s, which did not differ significantly from the latency of inhibitory responses in hippocampus (t_38_=0.0, p=1) or septum (t_54_=.053, p=.6) and also did not differ from the latency of excitatory responses in striatum (t_41_=1.62, p=.11).

#### 2.1.2 SWR responses in the bucket versus on the maze

SWR responses were analyzed separately in the bucket versus on the maze. Across all brain regions, n=52 neurons responded selectively to one behavioral condition but not the other (bucket but not maze, or vice versa), with the majority of these neurons (48/52, or 92.3%) located outside of the hippocampus. Of these behavior-selective neurons, 88.5% (46/52) responded only in the bucket but not on the maze, whereas 11.5% (6/52) responded on the maze but not in the bucket. Hence, neurons that responded to SWRs only in the bucket were much more prevalent than neurons that responded only on the maze (p<.0001 for binomial test against 50% probability for each preference). One possible explanation for this bias could be that rats spent more time sitting still in the bucket than on the maze during each session (Fig. 1B), and therefore, the statistical power for detecting SWR responses was generally higher in the bucket than on the maze. To control for such bias in statistical power, a downsample-and-shuffle approach was used to equalize the statistical power for detecting SWR responses on the maze and in the bucket (see Methods). After controlling for sampling bias in this way, the total number of behavior-selective cells in all brain regions fell from 52 to 32; among these, the proportion of cells that responded to SWRs only in the bucket but not on the maze was 87.5% (28/32), and the proportion of cells that responded to SWRs only on the maze but not in the bucket was 12.5% (4/32). Hence, controlling for SWR sampling bias did not change the proportion of behavior-selective cells that were responsive in the bucket versus on the maze, *χ*^2^ (1,*N*=84)=.018, p=.89. It should also be noted that the amplitude of SWR responses was larger in the bucket than on the maze (Fig. 1D), which could have influenced detection accuracy.

Across all SWR-excited and SWR-inhibited neurons combined, the proportion of neurons that responded selectively to SWRs under only one behavioral condition (either on the maze but not in the bucket, or vice versa) differed significantly among brain regions, *χ*^2^ (1,*N*=280)=61.0, p<.00001. A more detailed analysis of responding in the maze versus bucket for each brain region is presented below.

##### 2.1.2.1 Hippocampal Units

Fig. 3A shows that in the hippocampus, 97% (133/137) of SWR-excited and 100% (10/10) of SWR-inhibited neurons responded to SWRs both on the maze and in the bucket (see examples in Fig. 2F). For excited and inhibited neurons combined, 97.3% (143/147) of hippocampal units responded to SWRs both on the maze and in the bucket. Hence, hippocampal neurons showed very little selectively of their SWR responses for one behavioral condition over the other; almost all hippocampal neurons that responded to SWRs were responsive both in the bucket and on the maze.

##### 2.1.2.2 Septum Units

Fig. 3B shows that in septum, 71.9% (46/64) of SWR-excited neurons (examples shown in Fig. 4A) and 67.7% (21/31) of SWR-inhibited neurons (examples shown in Fig. 4D) responded to SWRs in both behavior conditions, whereas 23.4% (15/64) of SWR-excited neurons (examples shown in Fig. 4B) and 25.8% (8/31) of SWR-inhibited neurons (see example, Fig. 4E) responded to SWRs only in the bucket but not on the maze, and 4.7% (3/64) of SWR-excited neurons (see example, Fig. 4C) and 6.5% (2/31) of SWR-inhibited neurons responded to SWRs only on the maze but not in the bucket. The proportion of septum units exhibiting behavior-selective SWR responses did not differ significantly for SWR-excited versus SWR-inhibited neurons, *χ*^2^ (1,*N*=123)=.09, p=.76. Hence, excitatory and inhibitory responses to SWRs in septum were similarly dependent upon the rat’s behavioral state.

**Figure 4.**
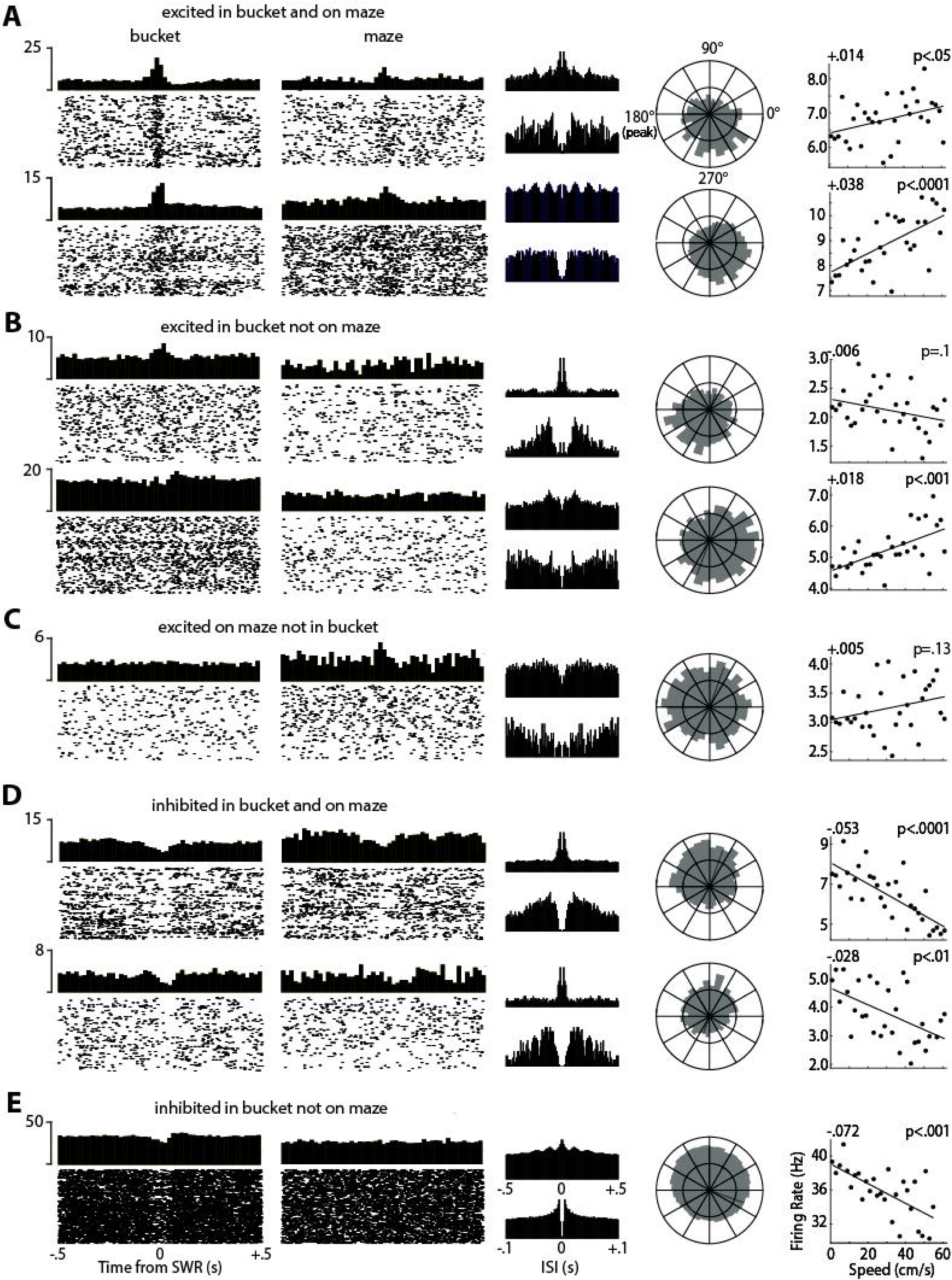
Example cells recorded in septum. Two left columns show PETHs and rastergrams aligned to SWR events that occurred in the bucket (first column) or on the maze (second column). Third column shows autocorrelograms of interspike intervals (ISIs) between -.5 and +5 s (top graph in each row) to illustrate theta rhythmicity, and between and -.1 and +.1 s (bottom graph in each row) to illustrate spike refractory periods. Fourth column shows circular distribution of spike phases relative to theta rhythm in the hippocampal LFP. Fifth column shows scatterplot in which each point is the cell’s mean firing rate (y-axis) at a given running speed (x-axis); regression line shows linear fit to the scatter points, with slope of speed modulation (in Hz/cm/s) at upper left and p-value of Pearson correlation at upper right. Examples are shown for septal neurons that were excited by SWRs in both the bucket and the maze **(A)**, excited by SWRs in the bucket but not the maze **(B)**, excited by SWRs on the maze but not in the bucket **(C)**, inhibited by SWRs in both the bucket and the maze **(D)**, and inhibited by SWRs in the bucket but not on the maze **(E)**.

##### 2.1.2.3 Striatal Units

Fig. 3C shows that in striatum, 42.9% (6/14) of SWR-excited neurons (examples shown in Fig. 5A) and 50% (12/24) of SWR-inhibited neurons (examples shown in Fig. 5C) responded to SWRs in both behavior conditions, whereas 50% (7/14) of SWR-excited neurons (examples shown in Fig. 5B) and 50% (12/24) of SWR-inhibited neurons (see example, Fig. 5D) responded to SWRs only in the bucket but not on the maze. The proportion of striatal units exhibiting behavior-selective SWR responses did not differ significantly for SWR-excited versus SWR-inhibited neurons, *χ*^2^ (1,*N*=58)=.06, p=.81. Hence, excitatory and inhibitory responses to SWRs in striatum were similarly dependent upon the rat’s behavioral state.

**Figure 5.**
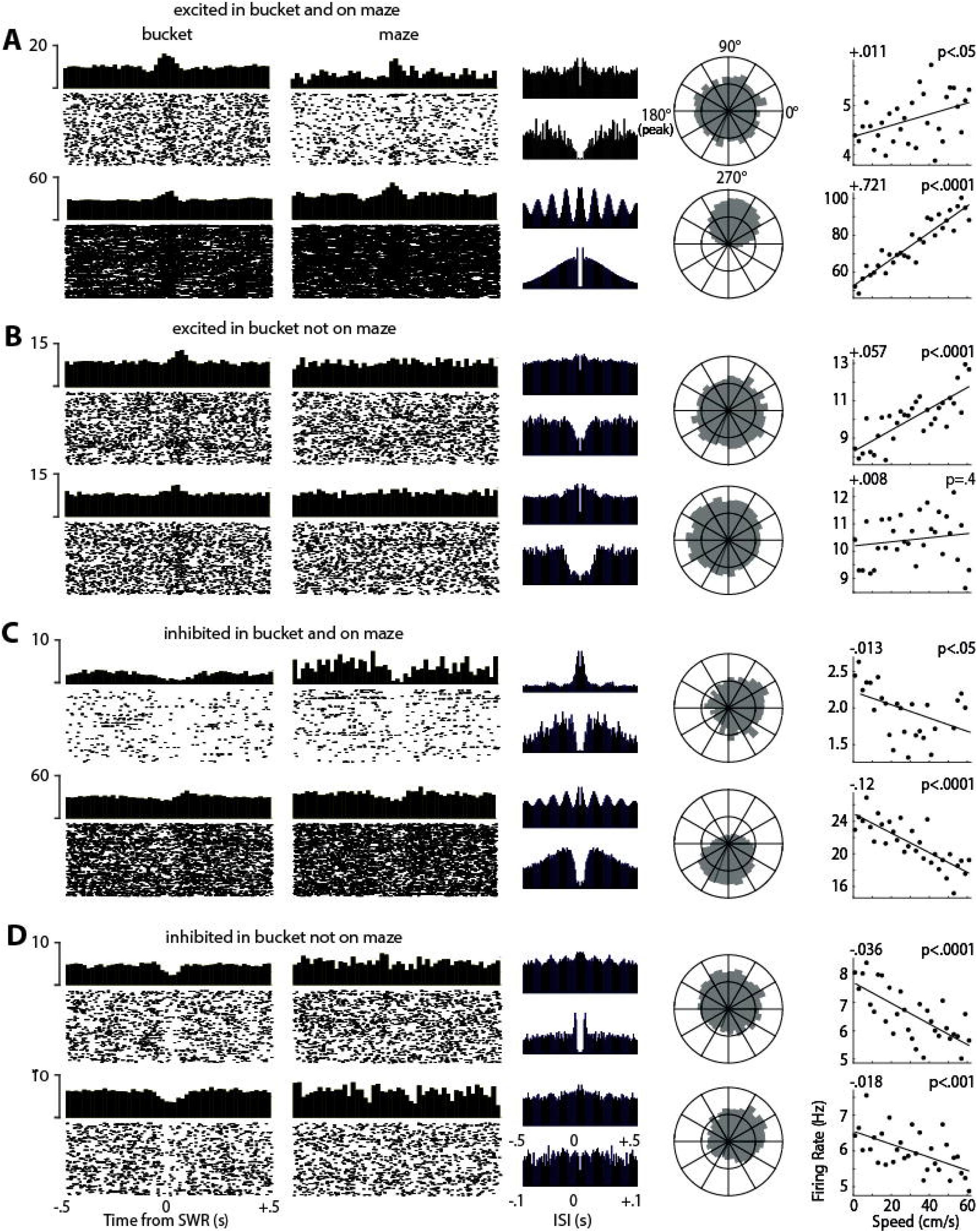
Example cells recorded in striatum. Graphs are the same as described in Figure 4. Examples are shown for striatal neurons that were excited by SWRs in both the bucket and the maze **(A)**, excited by SWRs in the bucket but not the maze **(B)**, inhibited by SWRs in both the bucket and the maze **(C)**, and inhibited by SWRs in the bucket but not on the maze **(D)**.

### 2.2 Predictors of Ripple Responsiveness

#### 2.2.1 Coherence with hippocampal theta rhythm

To quantify coherence of each neuron’s spike train with hippocampal theta rhythm, we measured the phase of theta rhythm at which individual spikes occurred, and then plotted the circular distribution of phases over all spikes generated by the neuron during active movement on the maze (running speed >10 cm/s). As in prior analyses (see above), we retain the convention that the valley and peak of theta rhythm occur at phases of 0° and 180°, respectively. For population analyses, the length of the phase distribution’s resultant vector (which ranged between 0 and 1) was taken as a measure of each cell’s spike coherence with hippocampal theta rhythm, where values near 0 indicate low coherence, and values near 1 indicate high coherence. It should be noted that coherence does not measure theta rhythmicity in a cell’s spike train, but instead measures the tendency of the cell to spike at a specific phase of the hippocampal LFP. That is, coherence measures periodicity in the cross-correlation between the spike train and the LFP, not in the auto-correlation of the spike train with itself, and it is possible to have one without the other (Zeitler et al., 2006). In some cases, neurons that exhibited strong spike coherence with hippocampal theta rhythm also exhibited strong theta rhythmicity of their own spike trains (for example, see the striatal unit in the bottom panel of Fig. 5A). But in other cases, neurons exhibited strong coherence with hippocampal theta rhythm while exhibiting very little evidence of theta rhythmicity their own spike train (for example, see the septal unit in the top panel of Fig. 4D).

To compare theta coherence of different neural populations, parametric statistical tests were performed upon the log_10_ of the resultant lengths (rather than raw resultant lengths), because log_10_ resultants were normally distributed whereas raw resultants were not. A 3-way independent ANOVA revealed that log_10_ resultant lengths differed significantly for neurons recorded in hippocampus, septum, and striatum (F_2,819_=71.47, p=2.5e-29). Post-hoc comparisons revealed that log_10_ resultants were not very different for hippocampal versus septal neurons (t_441_=2.16, p=.03; not significant at the .05 level after Bonferroni correction), reflecting the fact that both hippocampal and septal neurons were strongly coherent with hippocampal theta. However, striatal neurons exhibited much less theta coherence than either hippocampal neurons (t_591_=10.81, p=5.4e-5) or septal neurons (t_602_=8.75, p=2.1e-17).

For analyses of individual cells, a neuron was classified as “theta coherent” if a Rayleigh test for circular non-uniformity of its phase distribution yielded p<.01. The preferred firing phase for cells meeting this theta coherence criterion was then measured as the circular mean of the phase distribution (see below). This criterion classified 87% (188/216) of all hippocampal neurons as theta coherent, whereas 65.9% (149/226) of septal neurons and 25.1% (95/378) of striatal cells were classified as theta coherent. To further investigate whether any relationship existed between theta coherence and SWR responsiveness, additional analyses were carried out on cell populations from each area.

##### 2.2.1.1 Hippocampus

A 3-way independent ANOVA (Fig. 6, left) revealed that log_10_ resultant lengths differed for hippocampal neurons that were excited by, inhibited by, or non-responsive to SWR events (F_2,215_=3.08, p=.0478). Post-hoc comparisons indicated that SWR-excited cells exhibited significantly larger log_10_ resultants (and thus, more theta coherence) than non-SWR responsive cells (t_204_=2.4, p=.0174). By contrast, SWR-inhibited cells did not exhibit significantly different theta coherence from non-SWR responsive cells (t_77_=1.22, p=.22) or from SWR-excited cells (t_145_=.54, p=.58). However, the sample size of SWR-inhibited neurons was quite small (n=10), so statistical comparisons of these neurons against other populations may have been underpowered. Consistent with these population analysis results, Fig. 7A (left column) shows that the proportion of individual SWR-excited cells that were theta coherent was 92.7% (127/137), and the proportion of SWR-inhibited cells that were theta coherent was similarly high at 90% (9/10). By contrast, only 75.3% (52/69) of non-SWR responsive cells were theta-coherent. A 2×2 chi-square test (coherent vs non-coherent, SWR-responsive vs non-responsive) indicated that for septum cells, a significantly greater proportion of SWR-responsive than non-responsive cells were theta coherent, *χ*^2^(1,*N*=216)=12.24, p<.001.

**Figure 6.**
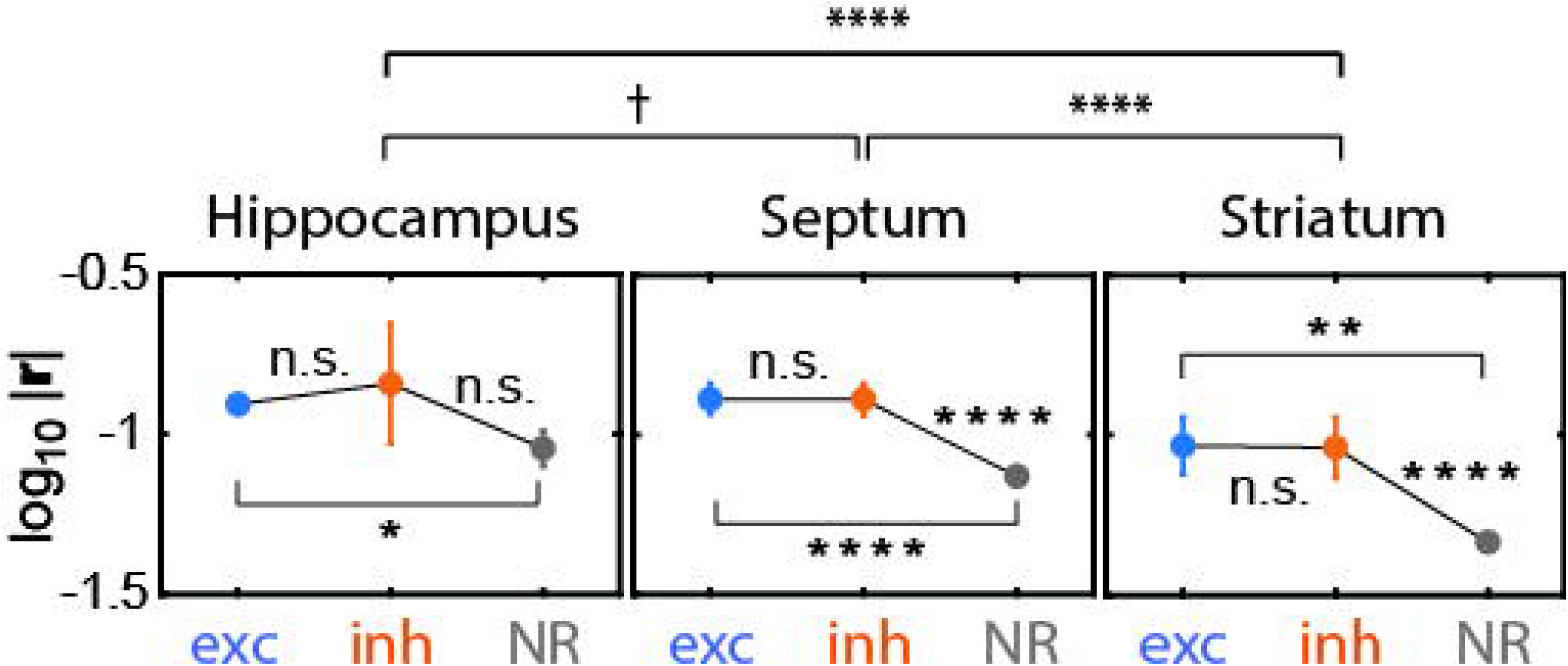
Theta coherence and SWR responsiveness. Graphs show mean log_10_ resultant lengths of preferred spike phases for cells that were excited (exc), inhibited (inh), or non-responsive (NR) to SWR events in hippocampus (left), septum (middle), and striatum (right). Symbols: □ p<.1, * p<.05, ** p<.01, **** p<.0001.

**Figure 7.**
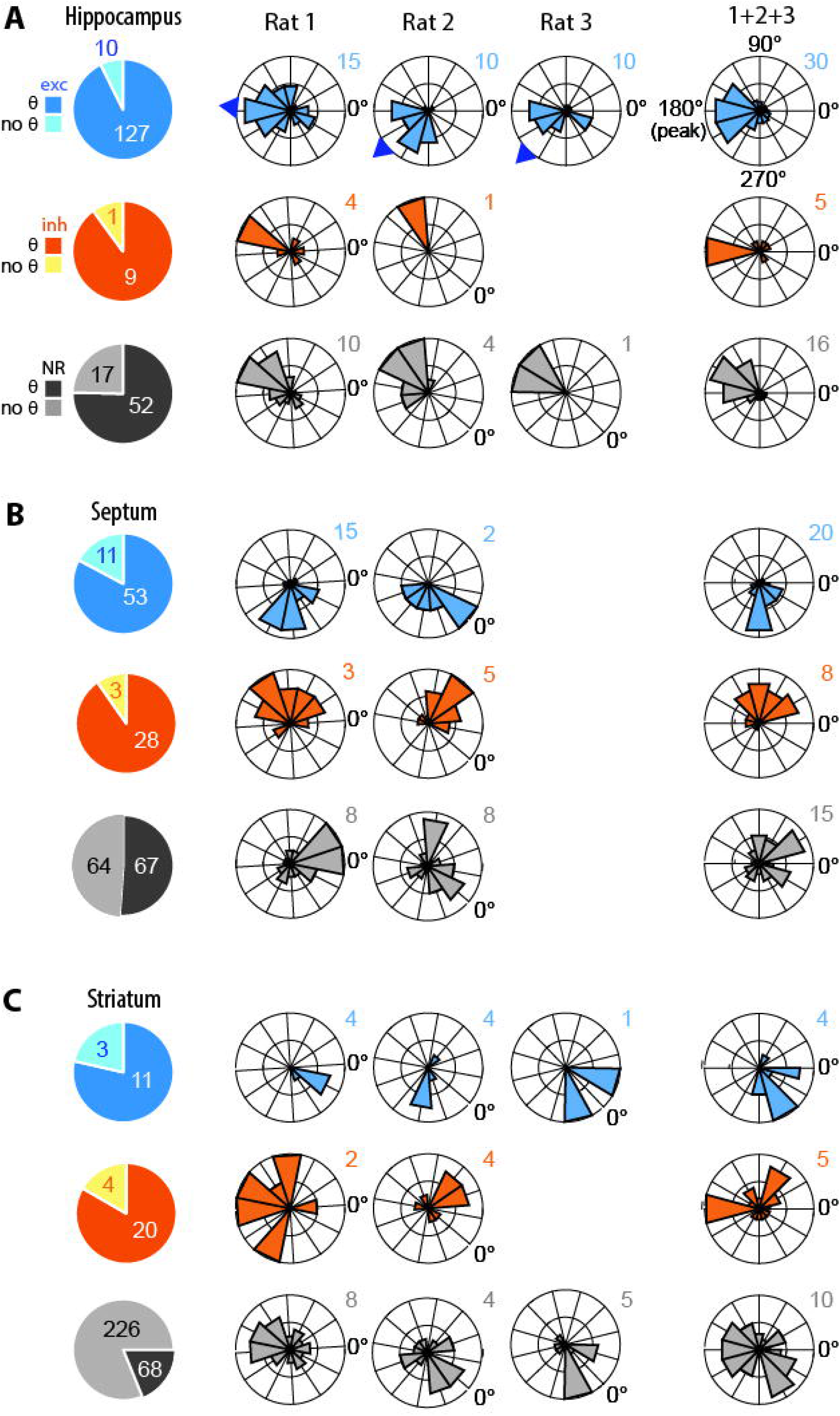
Preferred theta phase and SWR responsiveness. **A)** Pie charts at left show proportions of hippocampal neurons that were excited (exc), inhibited (inh), or non-responsive (NR) to SWR events that spiked coherently (□) or non-coherently (no □) with LFP theta rhythm. Polar plots at right show circular distributions of preferred firing phase for □-coherent cells in each individual rat (middle columns), and pooled across rats (right column). Arrows at rims of polar plots for individual rats in top row show mean theta phase of SWR-excited hippocampal cells in each rat. These means were used as a correction angle to align spike phases prior to averaging across rats (see main text). Below the top row, polar plots for individual rats in each column are rotated by the correction angle for that rat, to show how spike phases were aligned with one another prior to being pooled in the rightmost column. **B,C)** Same as ‘A’ for neurons in septum **(B)** and striatum **(C)**.

We next analyzed the phase of theta rhythm at which theta-coherent neurons fired within each individual rat. Rayleigh tests revealed that the preferred phases of theta coherent hippocampal neurons that were excited by SWRs (Fig. 7A, top row) were non-uniformly distributed in all three rats (Rat 1: Z_73_=6.3, p=.0017; Rat 2: Z_28_=16.6, p=2.8e-9; Rat 3: Z_26_=5.2, p=.0047). Moreover, SWR-excited cells had mean preferred phases concentrated near the peak of theta at 180° (mean phase for Rat 1: 176.4°, Rat 2: 220.4°, Rat 3: 226.1°). Circular V-tests revealed that the mean preferred phase of SWR-excited hippocampal neurons was not distinguishable from 180° for any of the rats (Rat 1: V_73_=21.4, p=2.0e-4; Rat 2: V_28_=16.4, p=5.7e-6, Rat 3: V_26_=8.0, p=.0128). Hence, SWR-excited hippocampal cells tended to fire near the peak of theta rhythm in all three rats. Theta coherent hippocampal cells that were inhibited by SWRs (Fig. 7A, middle row) were recorded from 2 of the 3 rats, and also exhibited mean preferred phases near the peak of theta rhythm (Rat 1: 154.8°, Rat 2: 147.7°). Sample sizes of SWR-inhibited hippocampal cells were too small in each rat to perform adequately powered within-animal Rayleigh or V tests (but see below for pooled analysis across all rats). Theta coherent hippocampal neurons that did not respond to SWRs were recorded from all three rats, and again exhibited mean preferred phases near the peak of theta rhythm (Rat 1: 151.2°, Rat 2: 181.5°, Rat 3: 195.7°). Circular statistics were only performed on non-responsive cell data from Rats 1 and 2, since only 2 non-responsive hippocampal cells were recorded from Rat 3. Preferred phases of non-responsive neurons were non-uniformly distributed in Rat 1 (Z_42_=8.42, p=1.5e-4) and Rat 2 (Z_16_=6.81, p=5.8e-4), and the mean cluster phase was not distinguishable from 180° in Rat 1 (V_37_=15.4, p=1.7e-4) or Rat 2 (V_13_=10.9, p=9.6e-6).

The above results from within-rat analyses indicated that most hippocampal neurons were theta coherent, and most tended to fire near the peak of theta rhythm (regardless of whether they responded to SWRs). There was some variability in the precise phase at which cells preferred to fire in different rats, and since the phase of the theta LFP is known to vary at different recording locations within the hippocampus (Lubenov & Siapas 2009; Patel et al., 2012), some of the between-rat variability in mean preferred firing phases may have resulted from differences in the precise locations at which theta LFP reference electrodes were positioned in each rat. To compensate for this, phase preference data was corrected to a new reference frame before being pooled across rats. The preferred phase of each neuron from a given rat was corrected by an angle equal to the mean phase at which SWR-excited hippocampal cells fired in that rat, plus 180° (to retain the convention that the peak of theta falls at 180°). In this corrected reference frame, cells that tended to fire in phase with SWR-excited hippocampal cells had a phase preference of 180°, and cells that tended to fire in antiphase with SWR-excited hippocampal cells had a phase preference of 0°. When pooled across rats, SWR-excited hippocampal neurons exhibited significant clustering of their corrected phases near 180° (V_127_=54.6, p=3.6e-12), but of course this is entirely expected, since the mean phase of SWR-excited cells was used to correct the phases in each rat. SWR-inhibited neurons also showed a trend to cluster near 180° (V_9_=3.1, p=.07), even though the phase of SWR-excited cells was used for correction. However, this analysis was underpowered because of the small sample size for SWR-inhibited hippocampal neurons (n=9). Theta coherent hippocampal neurons that were non-responsive to SWRs also exhibited significant clustering of their corrected phases near 180° (V_52_=26.1, p=1.5e-7), so these cells tended to fire near the same phase as SWR-excited cells.

In summary, the firing phases of hippocampal neurons tended to cluster near the peak of the LFP theta rhythm, regardless of how the cells responded to SWR events. Since more than half of hippocampal neurons in each rat were excited by SWRs, and since the preferred firing phase of SWR-excited neurons tended to cluster near a common LFP phase in each rat, we used the mean phase of hippocampal SWR-excited cells in each rat to correct the LFP phases of cells recorded in other areas (septum and striatum) into a common angular reference frame. In analyses presented below, we shall use the term “corrected phase” to refer to firing phases that have been shifted by a rat’s mean phase of SWR-excited hippocampal neurons, whereas “uncorrected phase” shall refer to the raw spike phase measured against each rat’s unshifted LFP.

##### 2.2.1.2 Lateral Septum

A 3-way independent ANOVA (Fig. 6, middle) revealed that log_10_ resultant lengths differed significantly for septal neurons that were excited by, inhibited by, or non-responsive to SWR events (F_2,225_=12.53, p=6.9e-6). Post-hoc comparisons revealed that non-SWR responsive cells exhibited significantly smaller log_10_ resultants (and thus, less theta coherence) than either SWR-excited cells (t_194_=4.31, p=2.6e-5) or SWR-inhibited cells (t_161_=3.49, p=6.2e-4), but SWR-excited cells did not exhibit significantly different resultant lengths from SWR-inhibited cells (t_93_=.02, p=.98). Hence, on average, SWR-excited and SWR-inhibited septal neurons both exhibited significantly greater theta coherence than non-SWR responsive septal neurons. Consistent with these population analysis results, Fig. 7B (left column) shows that the proportion of individual SWR-excited septal cells that were theta coherent was 82.8% (53/64), and the proportion of SWR-inhibited cells that were theta coherent was similarly high at 90.3% (28/31). By contrast, only 51.2% (67/131) of non-SWR responsive cells were theta-coherent. A 2×2 chi-square test on these proportions indicated that theta coherence of septum neurons was significantly contingent upon whether or not the cell was responsive to (that is, either excited or inhibited by) SWRs, *χ*^2^ (1,*N*=226)=35.9, p<.00001.

The uncorrected mean preferred phases of SWR-excited septal neurons (Fig. 7B, top row) in Rat 1 and Rat 2 were 273° and 316°, respectively. Rayleigh tests revealed that the preferred phases of theta coherent SWR-excited neurons were non-uniformly distributed in Rat 1 (Z_47_=20.9, p=5.4e-11) and Rat 2 (Z_6_=2.6, p=.064). When phase values from Rat 1 and Rat 2 were corrected by the mean phase of SWR-excited hippocampal cells in each rat (see Methods) and then pooled across rats, the mean phase at which SWR-excited septal neurons fired was 277°, which is +97° from the 180° peak of hippocampal theta. Circular V-tests revealed that in both individual rats, the mean preferred phase of SWR-excited septal neurons was not distinguishable from being shifted by +90° relative to the peak of hippocampal SWR-excited cells (Rat 1: V_47_=31.3, p=5.3e-11; Rat 2: V_6_=3.9, p=.0122), and this remained true when data was pooled across both rats (V_53_=35.1, p=4.8e-12).

The preferred phases of SWR-inhibited neurons (Fig. 7B, middle row) were also non-uniformly distributed in both rats (Rat 1: Z_13_=4.8, p=.006; Rat 2: Z_15_=-8.2, p=8.8e-5). The mean uncorrected phases of SWR-inhibited septal neurons in Rat 1 and Rat 2 were 92.8° and 95.3°, respectively. When phase data from Rat 1 and Rat 2 were corrected to the hippocampal reference frame and pooled together, the mean phase at which SWR-inhibited septal neurons fired was 70°, or −110° from the 180° peak of hippocampal theta. Circular V-tests revealed that in both individual rats, the mean preferred phase of SWR-inhibited septal neurons was not distinguishable from being shifted by −90° relative to the peak of hippocampal theta (Rat 1: V_13_=7.9, p=9.6e-4; Rat 2: V_15_=11.1, p=2.7e-5), and this remained true when data was pooled across both rats (V_28_=14.34, p=4.8e-5).

The preferred phases of neurons that were non-responsive to SWRs (Fig. 7B, bottom row) were non-uniformly distributed in Rat 1 (Z_35_=7.0, p=6.9e-4), and trended toward non-uniformity in Rat 2 (Z_33_=-2.4, p=.09). The mean uncorrected phases of non-SWR responsive septal neurons in Rat 1 and Rat 2 were 345.7° and 356.4°, respectively. When phase data from Rat 1 and Rat 2 were corrected into the hippocampal reference frame and pooled together, the mean phase at which non-responsive neurons fired was 355.9°, or +176° from the peak of hippocampal theta. Circular V-tests revealed that in both individual rats, the mean preferred phase of SWR-inhibited septal neurons was not distinguishable from being in antiphase with hippocampal theta (Rat 1: V_35_=15.1, p=.0025; Rat 2: V_33_=14.9, p=.0073), and the same was true of the pooled data (V_68_=24.5, p=1.3e-5).

In summary, a majority of all septal neurons were coherent with hippocampal theta rhythm. Similar to the hippocampus, about 80-90% of SWR-responsive cells were theta coherent, whereas only half of non-SWR responsive cells were theta coherent. But unlike in the hippocampus, SWR-excited versus inhibited neurons in septum tended to fire at distinct phases pf hippocampal theta. SWR-excited cells clustered at +90° and SWR-inhibited cells clustered at −90° from the peak of hippocampal theta. Theta coherent cells that did not respond to SWRs clustered near the 0° valley of hippocampal theta.

##### 2.2.1.3 Striatum

A 3-way ANOVA (Fig. 6, right) revealed that log_10_ resultant lengths differed significantly for striatal neurons that were excited by, inhibited by, or non-responsive to SWR events (F_2,376_=11.23, p=1.8e-5). Post-hoc comparisons revealed that non-SWR responsive striatal cells exhibited significantly smaller log_10_ resultants (and thus, less theta coherence) than either SWR-excited cells (t_351_=3.07, p=.0023) or SWR-inhibited cells (t_361_=3.79, p=1.7e-4), but SWR-excited cells did not exhibit significantly different resultant lengths from SWR-inhibited cells (t_36_=.04, p=.96). Hence, on average, SWR-excited and SWR-inhibited striatal neurons both exhibited significantly greater theta coherence than non-SWR responsive striatal neurons. Consistent with these population analysis results, we found that the proportion of individual SWR-excited cells (Fig. 7C, left column) that were theta coherent was 78.6% (11/14), and the proportion of SWR-inhibited cells that were theta coherent was similarly high at 83.3% (20/24). By contrast, only 18.9% (64/340) of non-SWR responsive cells were theta-coherent. A 2×2 chi-square test on these proportions indicated that theta coherence of striatal cells was significantly contingent upon whether or not the cell was responsive to SWRs, *χ*^2^(1,*N*=348)=63.3, p<.00001.

Fig. 7C (top row) shows that the mean uncorrected phases of SWR-excited striatal neurons were somewhat similar in all three rats: 311.0° (Rat 1), 313.4° (Rat 2), and 4.0° (Rat 3). The sample size of SWR-excited neurons in the striatum each rat was too small to perform within-rat analysis of preferred phase distributions, but when phase data from all three rats was corrected into the hippocampal reference frame and pooled together, SWR-excited striatal neurons exhibited significant non-uniformity (Z_11_=6.0, p=.001) with a mean phase of 304.9°. This suggests that these neurons may have preferred to fire just prior to the valley of hippocampal theta. SWR-inhibited striatal neurons (Fig. 7C, middle row) that were theta coherent were recorded in only two of the three rats. In Rat 1, the preferred phases of these neurons showed only a trend for non-uniformity of their circular distribution (Z_10_=2.5, p=.08), with a mean preferred phase of 150.8°. In Rat 2, the preferred phases beat significance for non-uniformity on the circle (Z_10_=3.2, p=.04), with a mean preferred phase of 85.6°. When phase data from both rats was shifted into the hippocampal reference frame and pooled together, SWR-inhibited septal neurons did not beat significance for non-uniformity of their preferred phases (Z_20_=1.3, p=.28). The mean uncorrected phases of theta coherent septal neurons that were not responsive to SWRs (Fig. 7C, bottom row) were 127.8° (Rat 1), 301.6° (Rat 2), and 291.7° (Rat 3). Preferred phases of non-responsive neurons were non-uniformly distributed in Rat 3 (Z_12_=4.0, p=.0154), but not in Rat 1 (Z_37_=2.3, p=.1) or Rat 2 (Z_15_=.82, p=.44). When phase data from all three rats was corrected into the hippocampal reference frame and pooled together, the preferred phases of non-SWR responsive striatal neurons did not beat significance for non-uniformity of their preferred phases (Z_64_=.60, p=.55).

In summary, a majority (about 80%) SWR-responsive striatal neurons spiked coherently with hippocampal theta rhythm, even though a minority (about one fourth) of all striatal neurons were theta coherent. Hence, theta coherence was significantly more common among SWR-responsive than non-responsive striatal neurons. SWR-excited striatal neurons showed a tendency to fire prior to the valley of hippocampal theta, but SWR-inhibited and non-SWR responsive striatal neurons did not exhibit significant tendencies to fire at a specific preferred phase of theta.

#### 2.2.2 Speed sensitivity

To analyze modulation of neural firing rates by running speed, we performed a linear regression analysis upon plots of firing rate versus running speed for each recorded cell (see Methods). The slope of the regression line (in units of Hz/cm/s) was taken as an estimate for the slope of speed modulation, and the intercept of the regression line (in units of Hz) was taken as an estimate for the cell’s mean firing rate during stillness.

##### 2.2.2.1 Hippocampus

Less than half of the neurons recorded in hippocampus (85/216, or 39.3%) met criterion for inclusion in the analysis of speed modulation. This was because many hippocampal neurons were spatially tuned (data not shown), and cells that fired selectively at a specific location often did not fire across a wide enough range of running speeds to meet criterion for inclusion in the speed analysis. We found that 65/85 (76.5%) of eligible hippocampal neurons exhibited a significant linear correlation (p<.05) of their firing rates with running speed, and of these, 62/65 (95%) were positively and 3/65 (5%) were negatively correlated with speed (Fig. 8A, left). When the sign of the SWR response was ignored, a 2×2 chi-square test found no contingency between sharp wave responsiveness (responsive vs non-responsive) and speed modulation (modulated vs non-modulated), *χ*^2^(1,*N*=85)=.47, p=.49. Hence, hippocampal cells that responded to SWRs were no more or less likely to be speed modulated than cells that did not respond to SWRs. Of the cells that were positively correlated with running speed, 39/65 (60%) were excited by SWRs and 1/65 (1.5%, a single cell) were inhibited by SWRs. For the subset of hippocampal cells that were both speed modulated and SWR-responsive (Fig. 8A, right), the mean slope of speed modulation was positive (.078±.014 Hz/cm/s) and significantly greater than zero (Z_39_=5.52, p=3.3e-8). In summary, a considerable majority of the hippocampal cells that were eligible for speed analysis had firing rates that were positively correlated with running speed, regardless of whether they were responsive to SWRs.

**Figure 8.**
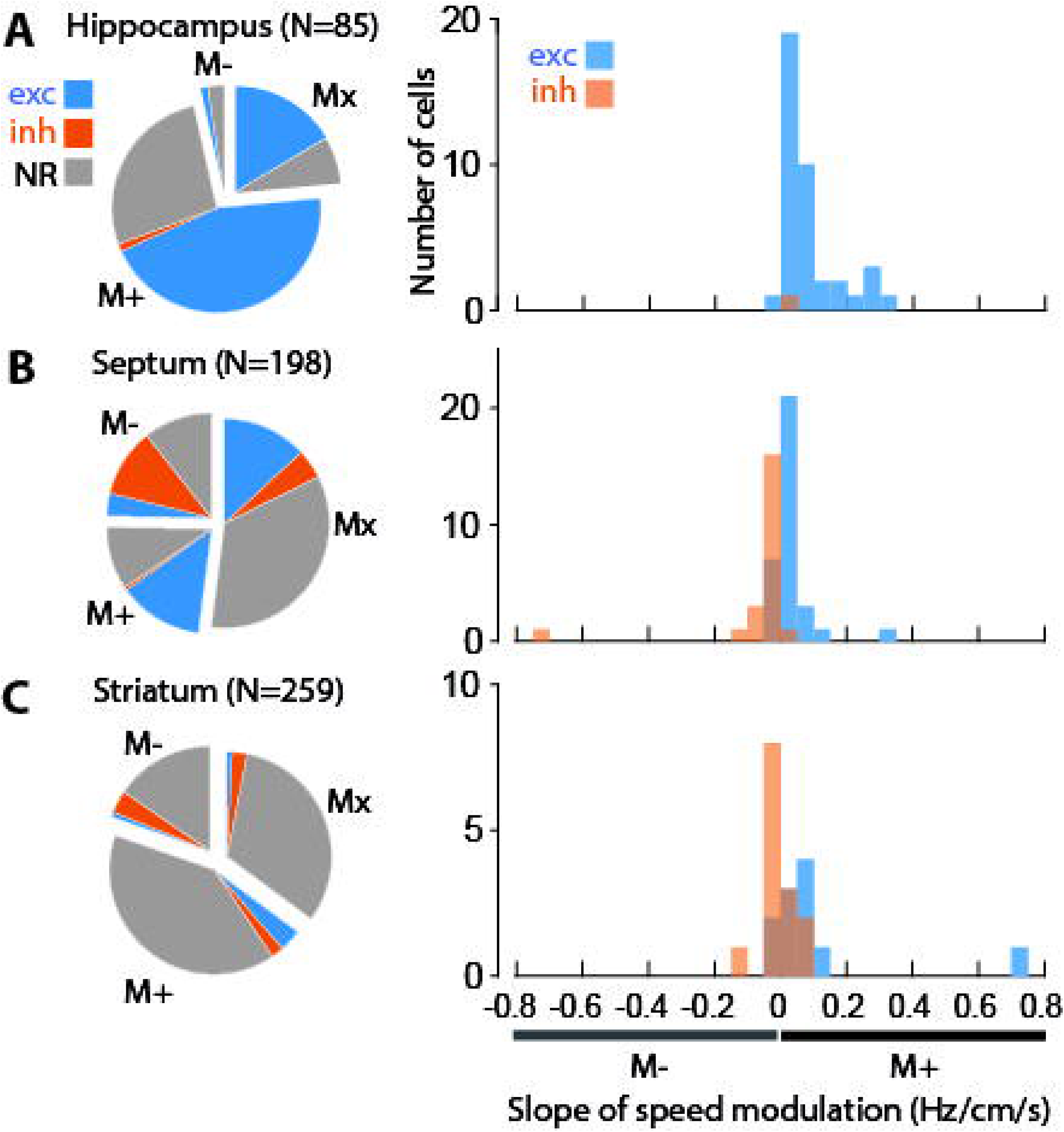
Speed modulation and SWR responsiveness. **A)** Pie chart shows proportions of hippocampal neurons that were eligible for speed analysis (N=85) that were positively (M+), negatively (M-), or not significantly (Mx) modulated by running speed. Within each speed classification, shading of wedges indicates proportions of cells that were excited (exc), inhibited (inh), or non-responsive (NR) to SWR events. Histogram at right shows the distribution of speed slopes for SWR-excited and SWR-inhibited cells that were significantly modulated by running speed. **B,C)** Same as ‘A’ for neurons in septum **(B)** and striatum **(C)**.

##### 2.2.2.2 Lateral Septum

Of the 226 neurons recorded in septum, 198 (87.6%) met criterion for inclusion in the analysis of speed modulation. We found that 95/198 (48%) of these neurons exhibited a significant linear correlation (p<.05) of their firing rates with running speed. Of the septal neurons that were speed modulated, 46/95 (48%) were positively and 49/95 (52%) were negatively correlated with running speed (Fig. 8B, left). Hence, speed-modulated neurons in septum were almost perfectly split in half between positively and negatively modulated cells, and this was true in both individual rats (rat #1: 26 positive, 25 negative; rat #2: 20 positive, 24 negative). When the sign of the SWR response was ignored, a 2×2 chi-square test found a significant contingency between sharp wave responsiveness (responsive vs non-responsive) and speed modulation (modulated vs non-modulated), *χ*^2^(1,*N*=198)=10.4, p=.0013. Hence, septal cells that responded to SWRs were more likely to be speed modulated than cells that did not respond to SWRs, in accordance with prior results (Wirtshafter & Wilson, 2019). A 3×2 chi-square test found a strong contingency between sharp wave responsiveness (excited, inhibited, or non-response to SWR) and speed modulation (positive vs negative), *χ*^2^(2,*N*=95)=29.2, p<.00001. When the sign of the SWR response was ignored, a 2×2 chi-square test found no contingency at all between sharp wave responsiveness (responsive vs non-responsive) and sign of speed modulation (positive vs negative), *χ*^2^(1,*N*=95)=.02, p=.88; hence, the sign of speed modulation did not depend upon whether or not a cell was responsive to SWRs. However, a chi-square test on just the SWR-responsive cells found that the sign of the SWR response was highly contingent upon the sign of speed modulation, *χ*^2^(1,N=55)=29.1, p<.00001. Indeed, only 1/27 (3.7%) of SWR-responsive septal cells with positive speed modulation slopes were inhibited by SWRs (the rest were excited by SWRs), and only 7/28 (25%) of SWR-responsive septal cells with negative speed modulation slopes were excited by SWRs (the rest were inhibited by SWRs). Consistent with this chi-square analysis, among cells that were both SWR responsive and speed modulated, the average slope of speed modulation for SWR-excited cells was significantly greater than zero (mean .028±.002 Hz/cm/s; Z_32_=2.41, p=.016), whereas the average slope for SWR-inhibited cells was significantly less than zero (mean -.06±.007 Hz/cm/s; Z_21_=-1.88, p=.06). Moreover, an independent t-test revealed that slopes of speed modulation for SWR-excited septal neurons were significantly more positive than slopes for SWR-inhibited septal neurons (t_53_=2.98, p=.0044). To make sure that this significant result did not arise solely from two outlying slopes with large values (see Fig. 8B, right), the t-test was re-run without these two outlying values, and despite a smaller difference between means, the result became even more statistically significant (t_51_=5.18, p=3.8e-6) because variance was reduced by eliminating the outliers.

These results show that among the subset of septal neurons that were both SWR responsive and speed modulated, cells with positive speed slopes were almost always excited by SWRs, and a considerable majority (about 75%) of cells with negative speed slopes were inhibited by SWRs. One possible confound is that this result might arise from a statistical power artifact, because SWRs occurred during stillness, and therefore, cells with negative speed slopes might tend to fire at a higher rates during stillness than cells with positive speed slopes. If so, then this may confer greater statistical power to detect SWR-induced inhibition of cells with negative speed slopes against their higher background firing rates during stillness, and greater statistical power to detect SWR-induced excitation of cells with positive speed slopes against their lower background firing rates during stillness. However, the median firing rate during stillness (estimated as the y-intercept of the speed slope line) did not differ (rank sum test Z=0.682, p=.5) for cells with positive (median 4.4 Hz) versus negative (median 3.8 Hz) speed slopes, nor did it differ (rank sum test Z=1.33, p=.18) for SWR-excited cells (median 4.6 Hz) versus SWR-inhibited cells (median 3.3 Hz). Hence, the correlation between the sign of the speed slope and the sign of the SWR response was unlikely to be a statistical power artifact.

##### 2.2.2.2 Striatum

Of the 378 neurons recorded in striatum, 259 (68.5%) met criterion for inclusion in the analysis of speed modulation. We found that 168/259 (64.8%) of these neurons exhibited a significant linear correlation (p<.05) of their firing rates with running speed, and of these, 117/168 (69.6%) had a positive and 51/168 (30.4%) had a negative slope of speed modulation (Fig 8C, left). Hence, in striatum, positively modulated cells were about twice as common as negatively modulated cells, although percentages varied somewhat across the three individual rats (rat #1: 32 positive, 33 negative; rat #2: 66 positive, 18 negative; rat #2: 19 positive, 0 negative). A 3×2 chi-square test found a significant contingency between sharp wave responsiveness and the sign of speed modulation for striatal neurons, *χ*^2^(2,*N*=168)=9.32, p=.0094. When the sign of the SWR response was ignored, a 2×2 chi-square test found only a weak trend for contingency between sharp wave responsiveness and sign of speed modulation, *χ*^2^(1,N=168)=2.31, p=.13; this trend indicated a modest tendency for negatively modulated speed cells to be more common among SWR-responsive than non-responsive cells in striatum. A chi-square test on just the SWR-responsive cells found that the sign of the SWR response was contingent upon the sign of speed modulation, *χ*^2^(1,N=26)=6.0, p=.014. Indeed, only 2/11 (18.2%) of SWR-responsive striatal cells with negative speed modulation slopes were excited by SWRs (the rest were inhibited by SWRs), and 5/15 (33.3%) of SWR-responsive cells with positive speed modulation slopes were inhibited by SWRs (the rest were excited by SWRs). Among cells that were both SWR responsive and speed modulated, the mean slope of speed modulation was positive for SWR-excited cells (.101±.019 Hz/cm/s), but not significantly greater than zero (Z_32_=1.59, p=.11). The mean slope of speed modulation was negative for SWR-inhibited cells (-.009±.003 Hz/cm/s), not significantly less than zero (Z_32_=-0.74, p=.45). However, an independent t-test revealed a trend for SWR-excited striatal neurons to have more positive speed slopes than SWR-inhibited neurons (t_23_=1.91, p=.069), and this effect reached significance (t_22_=2.63, p=.015) when variance was reduced by eliminating one extreme outlying slope which, despite being an outlier, did fit the overall trend for SWR-excited cells to have positive speed slopes (see Fig. 8C, right).

Taken together, these results show that among striatal neurons that were both SWR responsive and speed modulated, positively modulated speed cells tended to be excited by SWRs, and negatively modulated speed cells tended to be inhibited by SWRs. This correlation between speed slope and SWR responsiveness was less pronounced for striatal neurons than for septal neurons, but was nonetheless evident in both structures.

## 3. DISCUSSION

A growing body of evidence suggests that hippocampal projections to the septum may be an important route via which the hippocampus relays information to the midbrain and other subcortical regions to exert influence over behaviors such as reward-seeking, motor actions, reinforcement learning, and decision making (Luo et al., 2011; Gomperts et al., 2015; Tingley & Buzsaki 2018; Wirtshafter & Wilson 2019). Prior studies have demonstrated that septal neurons can encode an animal’s position in their firing rates (Takamura et al., 2006) as well as their spike phases (Tingley & Buzsaki 2018). Septal projections to the midbrain may thus relay position information from the hippocampus to dopaminergic and hypothalamic circuits that attach motivational value to specific spatial locations and environmental states (Luo et al., 2011; Gomperts et al., 2015; Wirtshafter & Wilson 2019; Tingley & Buzsaki 2020). Value and prediction error signals computed in the midbrain might then be relayed to cortical and striatal regions that govern learning, memory, and decision making, thereby allowing reinforcement learning processes be influenced by hippocampally encoded locations and states.

Compressed replay events that occur during SWRs have been hypothesized to play three distinct but related roles in reinforcement learning. First, it has been proposed that during navigation, forward replay of alternative future trajectories supports deliberation over the best path for the animal to take from its current location (Johnson and Redish, 2007; Pfeiffer & Foster, 2013; Yu & Frank, 2015; Wu et al., 2017; Kay et al., 2020). Second, it has been proposed that when reward outcomes are obtained, compressed replay of prior trajectories that have been traversed in the recent past may help to solve the “credit assignment” problem in reinforcement learning, which is the problem of assigning credit or blame for reward outcomes to decisions that were made in the remote past, before the outcome was obtained (Foster & Wilson, 2006). Third, it has been proposed that during sleep, compressed replay during SWRs may be necessary for consolidating short-term memories of recent experiences to long-term storage (Wilson & McNaughton, 1944; Buzsaki, 1996; Ego-Stengel & Wilson, 2010; Girardeau & Zugaro, 2011).

The septal output pathway from the hippocampus could play an important role in all three of these hypothesized functions for SWR events. Previous reports have demonstrated SWR-evoked responses in subpopulations of septal neurons (Wirtshafter & Wilson 2019; Tingley & Buzsaki 2020). It has also been shown that SWRs are sometimes accompanied by activation of midbrain neurons that respond to reward (Gomperts et al., 2015), as might be expected if dopamine circuits are computing value or prediction errors signals derived from replay of navigational trajectories during SWRs. Here, we recorded single units in the hippocampus, septum, and striatum while freely behaving rats ran trials in a T-maze task and rested in a holding bucket between trials. A large proportion of hippocampal neurons were excited during SWRs, as reported in prior studies (Wilson and McNaughton, 1994; Skaggs & McNaughton, 1996; Kudrimoti et al., 1999; Foster & Wilson, 2006; Davidson et al., 2009). We also identified several novel properties of SWR-responsive neurons in septum and striatum.

### 3.1 Activity of septal neurons during SWRs

In agreement with prior findings (Wirtshafter & Wilson 2019), we observed that SWR-responsive septal neurons tended to fire coherently with hippocampal theta rhythm during periods of locomotion. However, we also found that spikes of SWR-responsive septal neurons were segregated in time across the theta cycle, in such a way that SWR-excited neurons fired late in the cycle, whereas SWR-inhibited neurons fired early in the cycle. It has previously been shown that some septal neurons exhibit spatial phase precession against the hippocampal LFP as a rat runs on a maze (Tingley & Buzsaki 2018), but to our knowledge, it has not been previously reported that the valence (excitation versus inhibition) of a septal neuron’s SWR response during stillness is predictive of its preferred firing phase during theta rhythm when the animal is moving, as we found here.

In further agreement with prior findings (Wirtshafter & Wilson 2019), we observed that some SWR-responsive septal neurons behaved as speed cells, since their firing rates were positively or negatively modulated by the rat’s running speed. But interestingly, we also found (for the first time, as far as we know) that SWR-excited septal neurons tended to show positively sloped modulation of their firing rates by running speed, whereas SWR-inhibited septal neurons showed negatively sloped modulation of their firing rates by running speed. Taken together, our findings suggest there may be two distinct types of SWR-responsive neurons in septum: SWR-excited cells, which fire late in the hippocampal theta cycle and are biased to show positive modulation of their firing rates by running speed, and SWR-inhibited cells, which fire early in the theta cycle and are biased to show negative modulation of their firing rates by running speed.

It is interesting to consider how the spiking of SWR-excited versus inhibited septal neurons may align with the timing of place cell spikes in the hippocampus. As an animal passes through a place cell’s preferred firing location (or place field), the place cell bursts rhythmically at a slightly higher frequency than the LFP theta frequency, causing spikes to exhibit *phase precession* against the LFP (O’Keefe & Recce, 1993). At the population level, phase precession segregates place cell spikes in time, so that cells with place fields that lie ahead of the animal’s current location fire at late phases of LFP theta, whereas cells with place fields behind the animal’s current location fire early phases of LFP theta (Skaggs et al., 1996; Dragoi & Buzsaki, 2006; Wikenheiser & Redish 2013). Phase coding of spatial locations occurs in the lateral septum as well as the hippocampus (Tingley & Buzsaki, 2018; Monaco et al., 2019). Here, we found that SWR-excited septal neurons (which tend to be positively correlated with running speed) fired late in the theta cycle, so they presumably fired together with place cells that encoded locations ahead of the animal along its current motor trajectory. Conversely, we found that SWR-inhibited septal neurons (which tend to be negatively correlated with running speed) fired early in the theta cycle, so they presumably fired together with place cells that encoded locations behind of the animal along its current motor trajectory. Speed cells in the septum thus appear to have firing rates that are positively correlated with whichever behavior (running versus stopping) is most appropriate for reaching spatial locations that are encoded by place cells that co-fire with the speed cell during the theta cycle. This suggests that there may be a phase code for motor actions in septum that complements phase coding for position: speed cells that are positively correlated with movement (and excited during SWRs) may fire in phase with place cells whose preferred locations are reachable via continued movement, whereas speed cells that are negatively correlated with movement (and inhibited during SWRs) may fire in phase with place cells whose preferred locations are reachable via cessation of movement.

### 3.2 Activity of striatal neurons during SWRs

In the present study, neurons were recorded from both ventral and dorsal striatum. A prior study has shown that ventral striatal neurons exhibit phasic responses during dorsal hippocampal SWRs (Sosa et al., 2020), and we observed similar SWR responses in ventral striatal neurons. We also observed SWR responses in dorsal striatum, which to our knowledge has not been reported before. Previous single-unit recording studies have reported that firing rates of striatal neurons in rodents are correlated with the animal’s running speed (Ruede-Orozco & Robbe, 2015), and we similarly observed that a subset of SWR-responsive striatal neurons were modulated by running speed. Interestingly, SWR-excited striatal neurons tended to show positively sloped modulation of their firing rates by running speed, whereas SWR-inhibited striatal neurons showed negatively sloped modulation of their firing rates by running speed, similar to the results we obtained in septum.

Projection cells from the striatum are GABAergic medium spiny neurons (MSNs), which can be broadly subdivided into two main classes expressing D1 versus D2-type dopamine receptors. Classical models of the basal ganglia posit that D1 MSNs are the origin of a “direct” striatonigral motor output pathway which excites motor behavior, whereas D2 MSNs are the origin of an “indirect” striatopallidal motor output pathway which inhibits motor behavior. It would be worthwhile in future studies to investigate whether a significant proportion of SWR-responsive striatal neurons are MSNs, and if so, how excitatory versus inhibitory responses during SWRs are distributed among D1 versus D2 subtypes of MSNs. Neural recording and imaging studies have consistently failed to find evidence that D1 and D2 MSNs behave simply as motor-on and motor-off cells, as classical models would predict. Instead, both types of MSNs seem to fire together during initiation and execution of voluntary motor behaviors (Cui et al. 2013; Isomura et al, 2013), and combined with other evidence, these findings have led to speculation that D1 MSNs may help to drive the execution of selected actions, while D2 MSNs may simultaneously inhibit the execution of competing non-selected actions (Tecuapetla et al., 2016). It has been hypothesized that compressed replay by place cells during SWRs might provide a mechanism for animals to “deliberate” over decisions about which actions to select, and which actions to suppress (Yu & Frank, 2015). If so, then this could be regarded as tantamount to sorting out which actions should be excited by the D1 population and which should be suppressed by the D2 population during an impending motor decision. Consistent with this idea, it has been reported that reward-responsive midbrain dopamine neurons tend to fire synchronously with SWRs during wakeful stillness on a maze (but not during sleep), as might be expected if the animal were assessing the values of potential action plans during SWRs that occur on the maze (Gomperts et al., 2015).

Here, we recorded SWR events during stillness while overt motor actions were not being performed. Hence, one possibility to consider is that the SWR-responsive striatal cells we observed might be MSNs that are involved in inhibiting motor behavior, and thereby preventing actual motor actions from being performed during “virtual” navigation. Another possibility arises from prior evidence that acquisition of maze learning tasks is impaired by disruption of SWRs during both waking and sleep states (Girardeau et al., 2009; Jadhav et al., 2012), suggesting that SWRs may be directly involved in programming specific patterns of action selection that are required to achieve correct performance in such tasks. Since the striatum plays a key role in action selection and behavioral decision making, it could be that the SWR-evoked responses we observed in a small percentage of striatal neurons are reflective of a process by which MSNs become “programmed” to either excite or inhibit specific actions in the future, based upon value estimates for those actions that are generated during SWRs and compressed replay. This possibility could be further investigated in the future by experiments in which striatal unit activity is selectively disrupted during SWRs.

### 3.3 Summary and conclusions

SWRs are frequently accompanied by compressed replay of spatial trajectories within hippocampal place cell populations (Skaggs & McNaughton, 1996; Lee & Wilson, 2002; Foster & Wilson 2006; Diba & Buzsaki 2007; Davidson et al., 2009; Karlsson & Frank 2009). Findings presented here support the view that, in addition to being accompanied by hippocampal replay of spatial trajectories, SWR events might also be accompanied by activation of subcortical motor representations (Wirtshafter & Wilson 2019). Hence, when mental representations of a particular location become active within hippocampal place cell populations—either during an SWR event or during a “theta sequence” driven by phase precession—a corresponding representation of the motor action necessary to reach that location may become concurrently activated within subcortical regions, including septum and striatum. Concurrent activation of hippocampal state representations and subcortical action representations might support neural computations that are essential for reinforcement learning and value-based decision making.

Reinforcement learning theory (Sutton & Barto, 1998) suggests that value-based decision policies can be optimized by attaching values not just to particular states (such as residing at a specific spatial location) or particular actions (such as performing a specific motor behavior), but rather to *state-action pairs* (such as performing a specific action at a specific location). SWRs might therefore support reinforcement learning and decision making by activating representations of spatial trajectories and motor actions at the same time. For example, deliberation over alternative future trajectories during SWRs might not only involve activating hippocampal representations of spatial locations that lie along those trajectories (Johnson and Redish, 2007; Pfeiffer & Foster, 2013; Yu & Frank, 2015; Kay et al., 2020), but could additionally require activating representations of motor actions that must be performed at each location to adhere to a given trajectory. Similarly, when assigning credit for outcomes to recent behavioral choices (Foster & Wilson, 2006), re-activation of recently traversed trajectories during SWRs may require concurrent re-activation of the motor actions that were performed at each location along the trajectory. Finally, memory consolidation processes that require re-activation of recent experience during sleep (Wilson & McNaughton, 1944; Buzsaki, 1996; Ego-Stengel & Wilson, 2010; Girardeau & Zugaro, 2011) might necessitate concurrent reactivation of recently navigated spatial trajectories as well as motor actions performed along those trajectories, so that memories of decision policies can be consolidated by attaching values not just to states or to actions, but to state-action pairs that have previously yielded positive outcomes during waking experience.

## Acknowledgements

We thank Garrett Blair and Ryan Grgurich for valuable discussions. This work was supported by NSF NeuroNex grant 1707408 awarded to HTB (co-PI).

## 4. METHODS

All animal research protocols were reviewed and approved in advance by the UCLA Animal Research Committee, and conducted in accordance with United States federal guidelines. The data that support the findings of this study are openly available at DOI: 10.17632/rg3xjbgyjx.2.

### 4.1 Subjects and Behavior

#### 4.1.1 Subjects

Long-Evans rats (Charles River Laboratories, Hollister, CA, USA) were housed in a temperature and humidity controlled vivarium with a 12-12 reverse light-dark cycle, and fed ad lib until they attained a weight of ∼550 grams, after which they were reduced to 85% of their ad lib weight by limited daily feeding. The 3 rats used in the study were selected from a larger cohort of 6 rats that were all trained to perform a Figure 8 maze task prior to surgery. The three rats that were selected for surgery were the first three rats to reach a performance criterion (1 reward per minute over 20 minutes) on the Figure 8 maze.

#### 4.1.2 Behavior Apparatus

After recovery from surgery, rats were trained on a T-maze task. The three-arm T-maze was formed by blocking one of the arms on a four-arm plus maze apparatus. Throughout each block of trials, a barrier was placed at the entrance to one of the four arms, while the three remaining arms were assigned as the start, baited, and unbaited arms for the T-maze task. The maze was 218 cm wide with a 30 cm square platform in the center (see Fig. 1), located in a 3×3 m room with matte black walls and ceiling. Four 70 cm high posters with distinctive high-contrast black-and-white designs hung on the wall at the end of each arm to provide orienting landmark cues. The room was dimly lit by a 15 W light bulb aimed at the ceiling of the room. The reward was a ∼1 g piece of fresh banana. To make sure the rat was not guided by the strong odor of the banana, a dish containing a small amount of banana was always placed underneath the non-baited arm, inaccessible to the rat. Rats spent intertrial intervals in a holding bucket, from which they were not able to observe experimenters placing reward for the next trial.

#### 4.1.3 T-Maze Task

Rats were trained to run repeated acquisition and reversal trials on a T-maze (Fig. 1). At the start of each session, recording cables were connected and the rat was placed for 5 m in a white plastic bucket located next to the maze (the bucket always remained stationary in the same location, even as the start and goal arms were switched during different trial blocks) for a period of baseline recording. The rat was then placed by the experimenter at the designated start location for the current trial block, where it could immediately begin exploring the maze. The rat was free to run on the maze until it reached the end of either the baited or unbaited arm, at which point the experimenter placed the bucket behind the animal so that it the only available exit from the arm was to walk into the bucket. The rats usually climbed into and out of the bucket voluntarily, minimizing handling stress. The experimenter then placed the bucket in its assigned location on the floor beside the maze for a period of 1-3 m while the maze was cleaned and baited for the next trial. We cleaned the maze after each trial with 70% ethanol, and baited the reward arm for the next trial. When the rat completed 7/8 correct choice trials in a row, the baited and unbaited arms were swapped, and the rat began a reversal learning phase from the same start position. When the reversal criterion of 7/8 correct choice trials was reached, the barrier on the plus maze was moved to a different arm, so that the start, baited, and unbaited arms of the T-maze were reassigned. Another round of acquisition and reversal trials then began with the new maze configuration. This sequence of acquisition, reversal, and maze reconfiguration blocks continued throughout the entire duration of the recording experiment.

#### 4.1.4 Video tracking

The rat’s position was sampled at 30 Hz and tracked at a resolution of 2.2 pixels/cm by an overhead color video camera (JVC TK-C1480) outfitted with Tamron 2.8-12mm cctv CS aspherical lenses. The video signal was relayed to a position tracking system built into the electrophysiological data acquisition system (Neuralynx, Bozeman, MT). A custom offline algorithm compensated for lens distortion prior to analyzing the 2D position data.

### 4.2 Surgery, electrophysiology, and histology

#### 4.2.1 Surgery

Under deep isoflurane anesthesia, each rat was surgically implanted with a skull-mounted microdrive containing an array of 36 independently moveable probes. The 36 probes were grouped into 4 clusters, each consisting of 9 probes (8 tetrodes plus one reference) arranged in a diamond-shaped pattern where individual probes were spaced 400 μm from their nearest neighbors. Hence, the entire microdrive contained a total of 32 tetrodes and 4 reference wires. Of the 32 tetrodes, 16 were targeted at the dorsal hippocampus (8 per hemisphere), 6 were targeted at the lateral septum (3 per hemisphere), and 10 were targeted at the striatum (5 per hemisphere). In Rats 2 and 3, bilateral skull holes (each ∼2 mm in diameter) were centered at -/+3.2 ML and AP +1.1 (right) for dmStr/LS probe clusters, and at -/+3.4 ML and AP −4.5 for CA1 probe clusters. In Rat 1, skull holes were centered at -/+2.6 ML and AP +1.3 for dmStr/Nacc probe clusters, and at -/+2.0 ML and AP −2.8 for CA1 probe clusters. All coordinates are relative to Bregma. Rats recovered from surgery for at least 10 days before experiments began.

#### 4.2.2 Placement of LFP electrodes

After recovery from surgery, recordings were obtained while rats ran on the T-maze. On the first and second recording day, rats freely explored the maze for 15 minutes with no food rewards to acclimate to the environment. On the third day, rats began the initial acquisition phase of learning on the dual choice T-maze. During this initial training period, hippocampal tetrodes were advanced slowly into the CA1 layer of the hippocampus, until robust SWRs were detectable in the LFP on some of the tetrode wires, and robust 6-8 Hz theta rhythm was detectable on other tetrode wires. Data was not recorded from septal or striatal tetrodes during this initial training period (nor were the tetrodes advanced in these regions). When a hippocampal tetrode with robust SWRs was identified, and another tetrode wire with robust theta was found in the same hemisphere, these two tetrodes were chosen as the ripple and theta recording electrodes, respectively. Neither of these two tetrodes were advanced further during the remainder of the experiment. Starting with the next session, the goal and/or start arm was changed each time the rat achieved a criterion of 7/8 correct responses.

#### 4.2.3 Recording sessions and tetrode advancement

Throughout each maze session, a 128 channel DigitalLynx SX data acquisition system (Neuralynx, Bozeman, MT) was used to record LFP signals and single units at a sampling rate of 32 KHz per channel. LFP channels were high pass filtered above 1 Hz, and single-unit channels were bandpass filtered between 600-6000 Hz. Recording sessions varied in duration from 1 to 2 hours. At the end of a recording session, the rat occupied the bucket for 5 minutes before disconnection from the recording system. Tetrodes in septum and striatum were advanced by 333 μm after each session, so that different units would be recorded from these tetrodes in every session. By contrast, hippocampal tetrodes were advanced by at most 83 μm per day (and usually not at all), so that these tetrodes would remain within the hippocampal region throughout the entire experiment. Rats remained in the experiment area and rested in their home cages for at least 1 h before being returned to the vivarium for weighing and feeding.

#### 4.2.4 Histology

One day prior to euthanasia, the rat was deeply anesthetized with isoflurane and marking lesions were made on one tetrode wire per probe cluster by passing a 50 uA current through a lesion maker (Grass Instruments, West Warwick, RI) for 10 seconds at each polarity. 24 h after marking lesions were made, the rat was perfused transcardially with formalin and the fixed brain was carefully separated from the tetrode bundles, which were still positioned at their final advancement locations (we measured each probe’s linear excursion from the guide cannula to corroborate the advancement logs kept during the experiment). Brains were fixed in a solution of 30% sucrose formalin, sectioned at a thickness of 40 μm, and mounted on slides for imaging on a semi-automated digital light microscope (Keyence, Osaka, Japan). Slice images were referenced by overlaying them onto plates from the rat atlas of Paxinos & Watson (2004). Based upon marking lesions and track positions, the trajectory of each probe through the tissue was reconstructed by serial examination of all slices. The position of each tetrode on each recording day was estimated from the reconstructed trajectories.

### 4.3 Data analysis

#### 4.3.1 Spike Sorting

Manual spike sorting was performed offline using SpikeSort 3D (Neuralynx, Bozeman, MT). Cluster cutting was primarily performed based on the peak and valley amplitudes of spikes across all tetrode channels. In some cases, spike energy and PCA components 1, 2 and 3 were analyzed to achieve better separation. Clusters containing interspike intervals <1 ms were removed from analysis for lack of a refractory period.

#### 4.3.2 LFP filtering and analysis

On the assigned SWR probe for each animal, SWR events were detected as threshold crossings of the ripple band envelope which occurred when the rat was sitting still (movement speed < 2 cm/s). An 8^th^ order IIR filter was applied to extract signals in the 180-250 Hz band from LFP channel data sampled at 32 KHz. The envelope of the ripple band was taken as the absolute value of the Hilbert transform of the bandpass filtered signal. SWR events were detected as upward crossings of the ripple envelope amplitude past a threshold equal to 4 standard deviations above the mean envelope amplitude. The mean, standard deviation, and SWR threshold were calculated separately for data collected on the maze versus in the bucket, because SWR amplitudes differed for these two conditions (see Results). A lockout period of 100 ms was imposed after each SWR event, so that the next SWR event could not be detected until the lockout period had expired.

On the assigned theta LFP probe for each animal, a bidirectional 8^th^ order IIR filter was applied (using MATLAB’s ‘filtfilt’ command) to extract signals in the 4-12 Hz theta band from LFP channel data sampled at 32 KHz. Theta phase was derived using MATLAB’s ‘angle’ command from the Hilbert transform of the bandpass filtered signal.

#### 4.3.3 Response latency

To measure the latency between SWR events and a neuron’s spike responses, we computed a peristimulus histogram of spike responses (10 ms bins, spanning ±0.5 s) triggered at the peak of each SWR event’s ripple band LFP envelope. Two iterations of smoothing with a 50 ms (5 bin) boxcar window were performed, and the peak of the smoothed histogram was taken as the time of the peak unit response. As explained in Results, septal neurons and striatal neurons were never recorded during more than one session, but hippocampal neurons were often recorded over multiple sessions. In cases where a hippocampal neuron was recorded during more than one session, we identified the session for which the cell exhibited the most statistically significant (lowest p-value) spike response to SWRs, and used that session to measure the response latency.

#### 4.3.4 Unit responses during SWRs

To quantify a neuron’s response to SWR events, we counted the number of spikes that the neuron fired within a ±50 ms window surrounding each SWR, and divided by the width of the time window (100 ms) to compute the unit’s response rate (in Hz) during each SWR event. To measure the neuron’s baseline firing rate, we summed the number of spikes fired within two baseline windows on either side of the SWR event (−500 to −300 ms, and +300 to +500 ms), and divided by the summed width of both windows (400 ms) to compute a baseline firing rate (in Hz) for each SWR event. A neuron was considered to be responsive during SWRs if the response rate during SWRs was significantly different from the baseline response rate across all SWRs during which the neuron was recorded (see Results).

#### 4.3.5 Speed analysis

To analyze modulation of neural firing rates by running speed, position data from the video tracker (sampled at 30 Hz) was smoothed by convolution with a boxcar window 7 samples wide. Speed at each sample time *t* was then estimated at seven different lag times: L=33, 66, 99, 122, 155, 168, and 201 ms. The median of these seven estimates was then taken as the measure of speed at time *t*. The following formula estimated speed at sample time *t* using lag L:

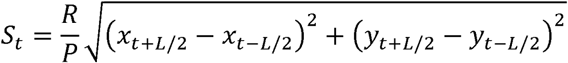

where *s*_*t*_ is the estimated speed, *R*=1000/*L* is the lag frequency in Hz, *P*=2.2 cm/pixel is the tracking resolution, and (*x*_*t*_, *y*_*t*_) is the interpolated position in pixels at time *t*. Linear interpolation of the speed time series (sampled at 30 Hz) was used to estimate the rat’s running speed at each spike time (sampled at 32 KHz). A cell’s firing rate at each running speed was computed by binning spike-triggered speed measurements in the range 0 to 60 cm/s using bins 2 cm/s wide, and then dividing the total number of spikes in each speed bin by the total time spent running at that speed. Linear regression then calculated the slope and intercept of the best linear fit to points on the speed curve. Bins containing <10 spikes or <2 s of occupancy time were omitted, and at least 4 valid bins were required for inclusion in the regression analysis.

